# Accurate, ultra-low coverage genome reconstruction and association studies in Hybrid Swarm mapping populations

**DOI:** 10.1101/671925

**Authors:** Cory A. Weller, Susanne Tilk, Subhash Rajpurohit, Alan O. Bergland

**Affiliations:** Department of Biology, University of Virginia, Charlottesville, Virginia 22904; National Human Genome Research Institute, National Institutes of Health, Bethesda, MD 20892; Department of Biology, Stanford University, Stanford CA 94305; Department of Biological and Life Sciences, Ahmedabad University, Ahmedabad, India 380009

## Abstract

Genetic association studies seek to uncover the link between genotype and phenotype, and often utilize inbred reference panels as a replicable source of genetic variation. However, inbred reference panels can differ substantially from wild populations in their genotypic distribution, patterns of linkage-disequilibrium, and nucleotide diversity. As a result, associations discovered using inbred reference panels may not reflect the genetic basis of phenotypic variation in natural populations. To address this problem, we evaluated a mapping population design where dozens to hundreds of inbred lines are outbred for few generations, which we call the Hybrid Swarm. The Hybrid Swarm approach has likely remained underutilized relative to pre-sequenced inbred lines due to the costs of genome-wide genotyping. To reduce sequencing costs and make the Hybrid Swarm approach feasible, we developed a computational pipeline that reconstructs accurate whole genomes from ultra-low-coverage (0.05X) sequence data in Hybrid Swarm populations derived from ancestors with phased haplotypes. We evaluate reconstructions using genetic variation from the Drosophila Genetic Reference Panel as well as variation from neutral simulations. We compared the power and precision of GWAS using the Hybrid Swarm, inbred lines, recombinant inbred lines, and highly outbred populations across a range of allele frequencies, effect sizes, and genetic architectures. Our simulations show that these different mapping panels vary in their power and precision, largely depending on the architecture of the trait. The Hybrid Swam and RILs outperform inbred lines for quantitative traits, but not for monogenic ones. Taken together, our results demonstrate the feasibility of the Hybrid Swarm as a cost-effective method of fine-scale genetic mapping.

## Introduction

Genetic mapping studies seek to describe the link between genotype and phenotype. For experimental crosses, mapping was traditionally conducted by scoring the phenotypes of recombinant offspring descended from a limited number of parental lines (Lander and Botstein 1989). While such QTL mapping studies can have high power to detect association, they offer minimal mapping resolution (Cheng *et al*. 2010), often detecting broad regions of phenotypic association. If linkage disequilibrium is lowered, regions of association can be resolved at the gene or nucleotide level (Li *et al*. 2005; Rockman and Kruglyak 2008), as in GWAS of large outbred populations (Nikpay *et al*. 2015; Wu *et al*. 2017; Monir and Zhu 2017). However, GWAS suffer from reduced power to detect associations (Long and Langley 1999), necessitating a large sample size relative to QTL mapping (Spencer *et al*. 2009).

To generate higher resolution mapping populations than the traditional biparental crosses, Multiparental Populations (MPPs) are becoming increasingly used. By crossing together multiple inbred lines, one can produce genetically diverse mapping populations without sampling wild individuals. MPPs are commonly used for the dissection of complex traits in model organisms (Chesler *et al*. 2008; Kover *et al*. 2009; King *et al*. 2012) and agriculturally important crops (Huang *et al*. 2012a; Singh *et al*. 2013; Krämer *et al*. 2014). Although MPPs are statistically powerful (Valdar *et al*. 2006; Svenson *et al*. 2012), they may have limited utility in addressing basic questions about the evolutionary forces affecting causal variants in natural populations (Long *et al*. 2014) and are often restricted to existing mapping panels in a limited number of taxa.

The mapping resolution of MPPs depends on the extent of linkage disequilibrium. Mapping resolution is improved by allowing for more recombination between haplotypes or by substituting extensive recombination for increased haplotype diversity (Mott *et al*. 2000; Chia *et al*. 2005). In the latter approach, by crossing dozens to hundreds of inbred lines for a limited number (~5) of generations and subsequently phenotyping and genotyping outbred individuals, heterozygous mapping populations can be generated quickly with sufficiently reduced LD to potentially detect associations with high resolution. We refer to such an outbred mapping population as a Hybrid Swarm. Whether Hybrid Swarm mapping populations can be generated efficiently, and whether association mapping using such a population is useful, remain open questions.

To address these questions, we evaluate a method to reconstruct phased whole genomes from large Hybrid Swarm populations using ultra-shallow sequencing (<0.05X) and assessed the power and precision of association mapping using a Hybrid Swarm relative to common alternatives [8-way MPP – DSPR (Long *et al*. 2014); inbred lines – DGRP (MacKay *et al*. 2012)]. First, we developed and tested our genome reconstruction method by generating whole genomes for thousands of simulated Hybrid Swarm individuals. Our simulated genomes draw from natural variation in the *Drosophila melanogaster* Genetic Reference Panel (DGRP), or variation generated from coalescent models representing a broad range of genetic diversity parameters for common model systems. We show that the Hybrid Swarm approach allows for highly accurate genotyping (average 99.9% genotypic accuracy) from ultra-low-coverage whole-genome individual-based sequencing. We confirm the accuracy of genome reconstructions using experimental animals from a Hybrid Swarm constructed in the lab from 128 DGRP founder lines. We then simulated a range of genetic architectures to discern the accuracy and precision of association mapping in the Hybrid Swarm compared to inbred lines, recombinant inbred lines, and a highly outbred population. GWA simulations confirm that inbred MPPs have the highest power to detect associations, outbred Hybrid Swarm populations have intermediate power, and inbred reference panels have the lowest power, particularly for quantitative traits. In addition, we show that outbred Hybrid Swarms eliminate spurious signals of association that arise using inbred lines. Together, our results show the feasibility of cost-effective association mapping in a large outbred multi-parental population and provides tools for genome-reconstruction and simulation.

## Methods

### Simulating a Hybrid Swarm

As a case study of low-coverage genome reconstruction in a model system, we simulated a Hybrid Swarm using sequenced inbred lines from the Drosophila Genetic Reference Panel (MacKay *et al*. 2012) as available from the *Drosophila* Genome Nexus (Lack *et al*. 2015). We only included the 128 lines (out of 205) with the least amount of missing genotype data. We removed insertions, deletions, and sites with more than two alleles. Any heterozygous genotype calls were masked as missing data.

To generate simulated populations, we developed a forward-simulator in R that stores ancestral haplotype block maps instead of genotypes. See Figure 1 for depictions of mapping populations simulated. We simulated Hybrid Swarms through random mating over five non-overlapping generations at a population size of 10,000. Sexual reproduction was simulated by random sampling of recombinant gametes from male-female pairs. Recombination frequency was modeled as a Poisson process with an expected value λ = Σ(Morgans) per chromosome. For simulations of *Drosophila* populations based on DGRP chromosomes, recombination occurred only in females, with recombination frequency and position based on values from Comeron et al (2012). See extended methods and Supplemental Figure S1 for additional detail.

**Figure 1.**
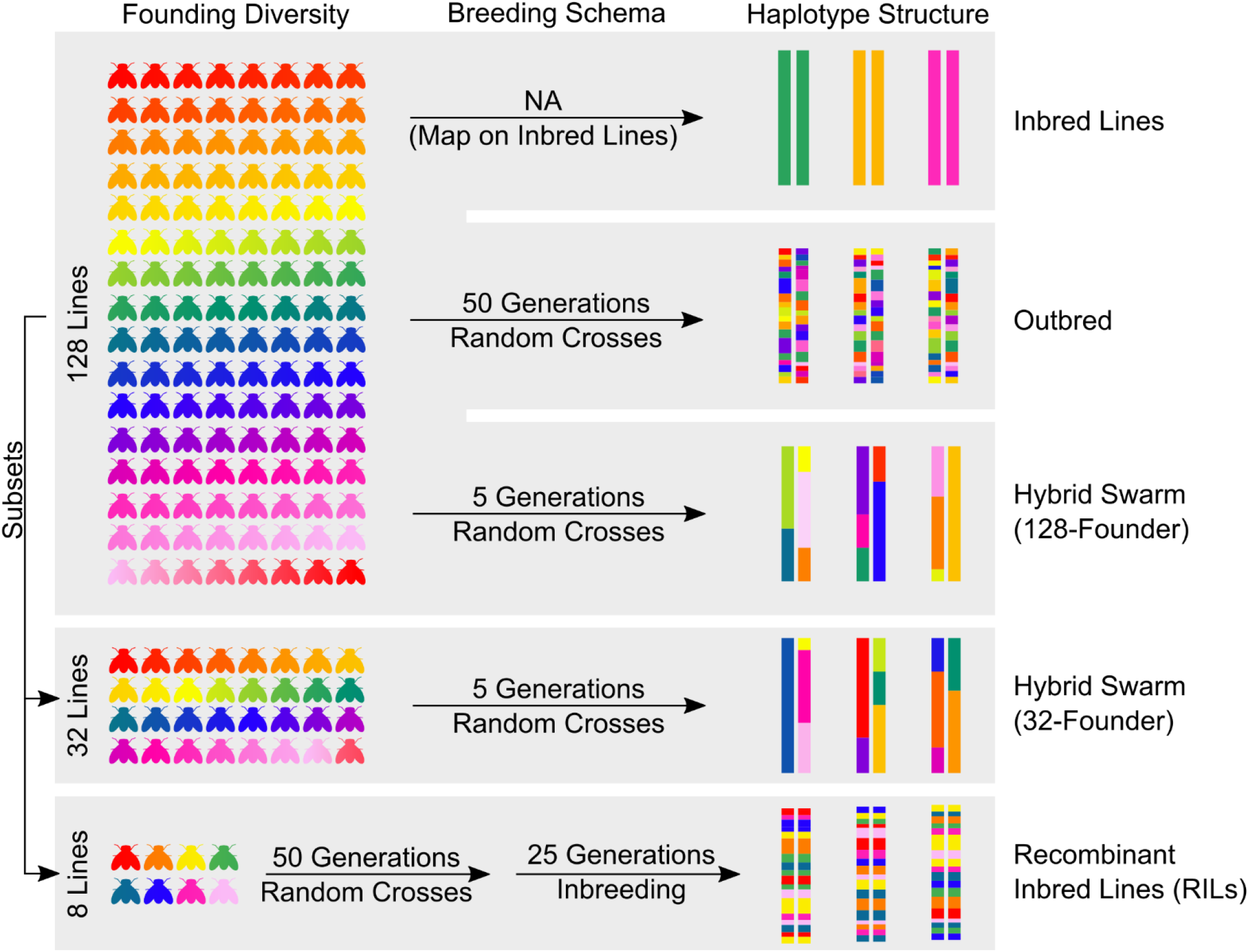
Depictions of various mapping populations explored. Each color represents a unique haplotype.

We also simulated Hybrid Swarms using haplotypes derived from whole-genome coalescent simulation using *scrm* (Staab *et al*. 2015). We generated simulations across a range of diversity and recombination levels, encompassing the values for many common model systems. For populations founded by simulated haplotypes, recombination occurred in both sexes, with recombination occurring uniformly across each chromosome.

We used *wgsim* (Li 2011) to simulate 100 bp reads at two average read depths (0.005X and 0.05X). We specified a base error rate of 0.001 and an indel fraction of 0. Remaining *wgsim* parameters were left as default. We assembled paired end reads using *PEAR* (Zhang *et al*. 2014) and separately aligned the assembled and unassembled groups to a reference genome with *bwa* 0.7.14 using the BWA-MEM algorithm (Li 2013). Simulated *Drosophila* reads were mapped to the *D. melanogaster* reference genome v5.39. After converting mapped reads to compressed BAM format with *samtools* 1.3.1 (Li *et al*. 2009), we removed PCR duplicates with *Picard tools* 2.0.1 (“Picard toolkit” 2019).

### Genome reconstruction for a Hybrid Swarm

Genome reconstruction for Hybrid Swarm individuals can take advantage of Hidden Markov Models developed for standard MPPs (Zheng *et al*. 2015b). In these models, the probability of each diplotype – a unique diploid combination of founding haplotypes - is calculated. Calculating these probabilities is feasible with smaller numbers of founding lines (< 8, typically) but grows at a quadratic rate. Therefore, increasing the number of founding haplotypes from 8 to 128 incurs orders of magnitude more computational effort (Supplemental Figure S2) and is not feasible. To make chromosome reconstructions in the Hybrid Swarm computationally tractable, we developed a method of accurately selecting a subset of most likely ancestors for any single chromosome. It is important to note that the computational requirement of selecting a subset of founding chromosomes requires that a Hybrid Swarm only be propagated for a limited number of generations, especially if the number of founding lines is large.

We used the software package *HARP* (Kessner *et al*. 2013) to identify the most likely founding lines based on the number of genomic windows with high probabilities of ancestry. *HARP* was originally developed to estimate haplotype frequencies from pooled sequence data, and we co-opted it to assess relative likelihood that any founder contributed to a genomic window in a focal Hybrid Swarm chromosome. We ran *HARP* with non-overlapping 100 kb windows with a minimum frequency cutoff 0.0001. Ideally, any given window would have four (or fewer) founders with high proportions of ancestry (~25%) and unambiguous haplotype paths (Supplemental Figure S3). However, empirical evaluation of Harp output for sample chromosomes showed that many had more than four founders with high proportion of ancestry and difficult-to-resolve haplotypes (Supplemental Figure S4).

We developed a heuristic method to identify the set of most likely ancestors. We ranked all possible founders by their contribution to the focal chromosome. To do this, we counted the number of windows with extreme likelihoods of ancestry (i.e., greater than a specified quantile) for each potential founding haplotype. Potential founding lines were then ranked by the number of windows passing this threshold. We examined two measures of effectiveness for this method across a range of quantile threshold values (90%, 95%, 99%, and 99.9%) when selecting up to a maximum number of most likely ancestral founders. The first measure is the number of true ancestral founders excluded; the second measure is the fraction of the chromosome derived from ancestors missing from the selected subset.

We used the Mathematica package RABBIT (Zheng *et al*. 2015a) to perform chromosome reconstructions. This package has been shown to be accurate for genotype estimation at sequencing coverage at 0.05X for a variety of multiparent populations (Zheng *et al*. 2018) but, the best of our knowledge, only for MPPs with twelve or fewer potential founders. To input the observed genotype of each recombinant individual, we counted reference and alternate reads at variable sites (i.e. polymorphic among the list of potential founders) using the Genome Analysis Toolkit *ASEReadCounter* tool (Broad Institute 2015). Because it is not possible to make confident homozygote genotype calls from low coverage sequencing data where most sites are observed only once and or twice, we encoded individuals with only reference or alternate alleles as “1N” or “2N”; observed heterozygotes were encoded as “12”. We sampled 5000 highly informative SNPs per individual per chromosome for genome reconstruction. Informative SNPs were identified as those at low frequency amongst potential founders identified in the HARP step. We ran RABBIT independently for each chromosome using the Viterbi decoding function under the “joint model” with all other RABBIT parameters left at default. RABBIT output was converted to a phased chromosome haplotype map, which we then used to extract and concatenate genotype information from a VCF file containing founder genotypes.

We evaluated the accuracy of reconstruction by calculating genotype accuracy and the number of inferred recombination events. To calculate genotype accuracy, we measured the fraction of sites where the estimated diploid genotype is identical to the originally simulated diploid genotype. We only examined accuracy on the autosomes. To measure accuracy of estimated frequency of recombination events, true and estimated recombination counts were first summed over both copies of each chromosome in a simulated individual. We then calculated Lin’s concordance correlation coefficient ρ between the true and estimated recombination counts using the *epi.ccc* function of the *R* package *epiR* (Stevenson 2018). An overview of our reconstruction pipeline is shown in Supplemental Figure S5.

### Chromosome reconstruction with STITCH

We estimated genotypes with alternative approach, STITCH, which is capable of imputing genotypes without reference panels (Davies *et al*. 2016). To assess genotype estimation accuracy using STITCH, we generated and mapped simulated reads for chromosome 2L at 0.05X coverage for 5000 individuals in a F5 Hybrid Swarm population (N=10000 individuals per generation). Because STITCH draws inference from haplotypes between related individuals, genotype estimate accuracy is expected to increase with greater numbers of sequenced individuals. To capture the effect of sample size on STITCH accuracy, we ran STITCH for sample sizes from 100 to 5000 sequenced individuals. We calculated genotype estimate accuracy as the fraction of variable sites with a correct diploid genotype estimate using a custom R script. Because STITCH memory requirements increase with greater values of *k* (founding haplotypes), it was not possible to evaluate *k*=128 founders, or to use the *diploid* estimation model. As a result, we limited our evaluations to 32-founder populations, using the *pseudoHaploid* model.

### Generating a real Hybrid Swarm population

We generated a hybrid swarm mapping population through undirected mating of all Drosophila Genetic Reference Panel (DGRP) inbred lines (Mackay et al. 2012) in large population cages (6’ x 6’ x 6’). Populations were seeded with males and females from each line and were expanded for four generations on cornmeal-molasses media. Cages reached an approximate final population size of ~500,000 as determined by a volumetric assay of dead adults. Generations were discrete: we removed egg-laden media from the cages and replaced egg-laden media in emptied cages after removal of the previous generation.

### Library preparation and read mapping

We homogenized individual flies in 350 μL Lysis Buffer RLT Plus using 2-3 1 mm beads in a bead shaker for 2 minutes. DNA and RNA were extracted using the AllPrep DNA/RNA Micro Kit (Qiagen product number 80284). Following DNA extraction, we purified samples with Ampure XP Beads (Beckman Coulter product number A63880). We prepared 1 μL of DNA at ~2.5 ng/μL sequencing using a modified Nextera protocol developed by Baym *et al*. (2015), indexing samples with custom primers, and selecting fragments between 450-500 bp in length using a SizeSelect e-Gel. As a final step to amplify the prepared DNA sequencing libraries, we ran all size selected samples through additional 5 rounds of PCR. Each PCR reaction used 5 μL template DNA, 0.6 μL of 100 mM forward and reverse primers (custom synthesized by IDT), 10μL of KAPA HiFi Ready Mix (KAPA Biosystem product number KK2611/2612), and 3.8 μL nuclease free water. Our PCR protocol included 5 minutes of initial denaturation at 95°C followed by 4 rounds of 20 seconds denaturation (98°C), 20 seconds annealing (62°C), 30 seconds elongation (72°C), followed by a final elongation at 72°C for 2 minutes. Following PCR amplification, we purified DNA libraries using Ampure XP beads and quantified concentrations on a Life Technologies Qubit spectrophotometer and Agilent Bioanalyzer. We sequenced the same library preparations at low (~0.05X) and high (~8) coverage on separate Illumina sequencing runs. Reads were mapped to the reference genome using the same methods as our simulations.

### Empirical evaluation of genotype accuracy in the Hybrid Swarm

We tested if RABBIT genotype estimates accurately reflected genotypes from higher coverage data for six individuals. First, we called genotypes in the high coverage samples using a heuristic based on reference and alternate allele counts produced by GATK’s ASEReadCounter to call genotypes. Sites with at least three reference and three alternate allele bearing short-reads were called as heterozygous; any site with at least six reference reads and zero alternate reads (or the converse) was called as homozygous. All other sites with fewer six reads were excluded. We calculated accuracy as the proportion of sites where the low-coverage genotype derived from reconstruction matched our higher-coverage genotype call. We note that our measurement of genotype accuracy from the empirical data is conservative as we expect a moderate fraction of true heterozygous sites to be called as homozygous.

### Simulating mapping populations for GWAS

We performed GWAS on simulations of various types of Hybrid Swarm designs and contrasted the signal of association using these designs to that from a simulated 8-way MPP (akin to the DSPR), and the DGRP. Simulation of GWAS proceeded in two steps.

For every type of genetic architecture or mapping population explored, we simulated 500 mapping populations comprised of 5000 individuals. Full genomes for individuals were generated using the forward simulation framework described above. We used the DGRP as the founder chromosomes. One important note is that the genotypes we use for GWAS are the “true” genotypes and are not affected by genotyping error. This decision was made to reduce computation time. Because genotyping error is randomly distributed throughout the genome (and rare on a per-site basis, see Figure 2), the use of perfect genotype data should not bias the results of our simulations.

**Figure 2.**
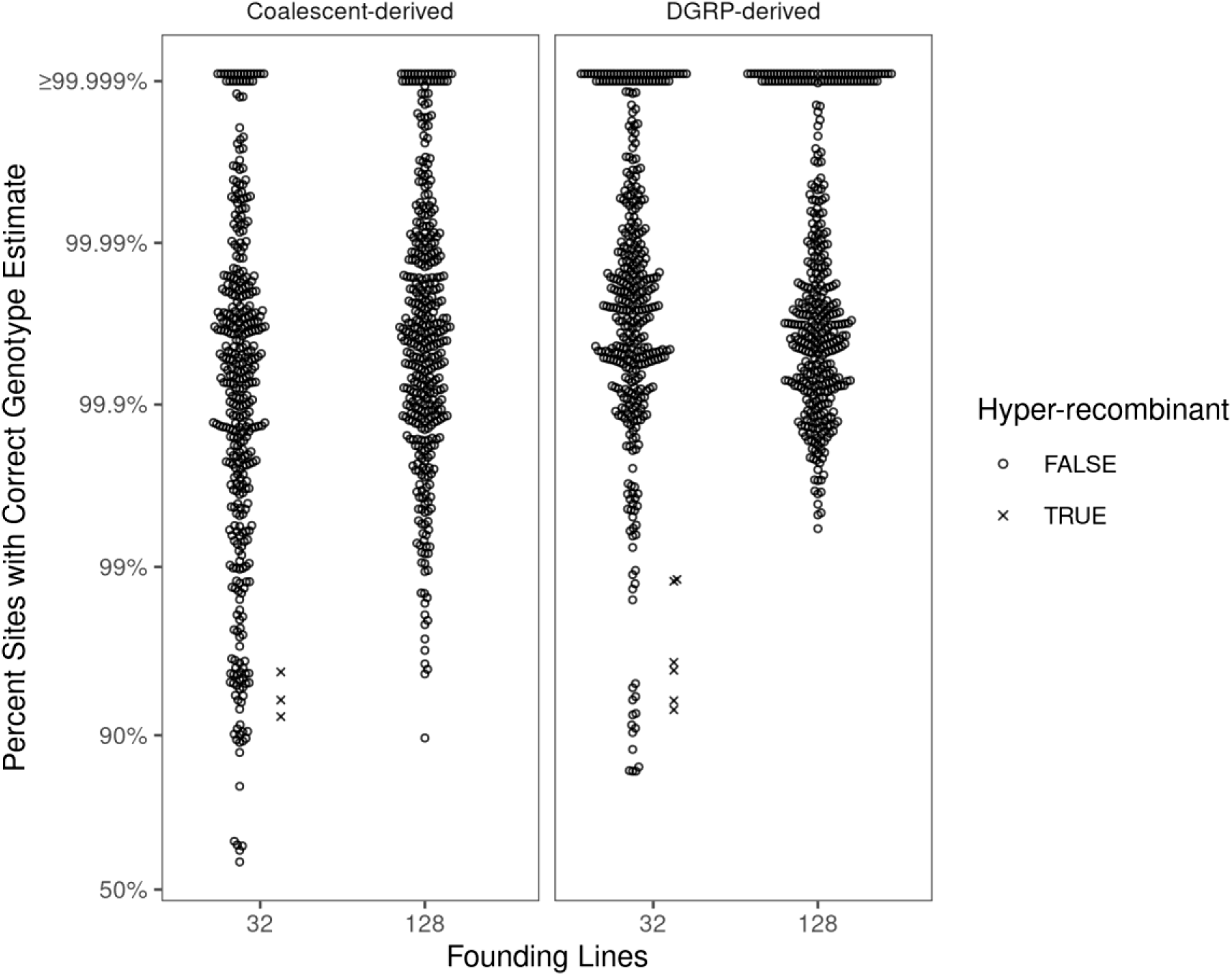
Genotype estimation accuracy for Hybrid Swarm populations. Reconstructions were performed for populations simulated using 32 or 128 inbred founder. Accuracy, calculated as the per-chromosome fraction of variable sites with a correct diploid genotype estimate, is shown on logit-transformed scale. Values are coded depending on the number of estimated recombination events, with highly recombinant estimates (>10 recombination events) displayed as an ×. Each parameter combination includes 400 reconstructed autosomes (individual circles) for 100 simulated individuals. The coalescent-derived individuals displayed here were simulated with an effective population size of N_e_=1 ×10^6^ and mutation rate μ=5×10^-9^.

Next, we assigned phenotypes to each simulated individual. We assign case or control phenotypes to individuals based on a liability model, where an observable binary phenotype is produced from an underlying continuous trait, known as its liability (Xu and Atchley 1996). Under this model, phenotypic variance will increase with additional causal loci. We contrasted mapping results for a monogenic trait and a polygenic trait, using loci with large and small effects, assigning individuals to case or control groups based on their genotype at causal loci. We randomly selected a causal locus (or multiple loci for the polygenic model) from all variable sites in the mapping population. We assigned the effect size of this locus to be proportional to its frequency: low frequency variants had a large effect and common variants had a small effect (Supplemental Figure S6). To achieve a population size of 5000, it is necessary to sample from the 128 inbred lines or 800 recombinant inbred lines with replacement.

To test for association between phenotype and each SNP, we performed a χ^2^ test of independence for reference and alternate allele counts between case and control groups. For inbred mapping populations, we corrected for non-independent allele draws by dividing the χ^2^ value by two.

### Assessing GWAS accuracy

To measure GWAS accuracy, we generated Receiver Operator Characteristic (ROC) curves and calculated Area Under the Curve (AUC). Briefly, we calculated the true positive rate (sensitivity) as a function of false positive rate (1-specificty) using the *R* package *pROC* (Turck *et al*. 2011). In our simulations, only the causal SNP may be considered a true positive. We summarize GWAS accuracy by averaging ROC curves from all replicate simulations, generating a single representative ROC curve following Marigorta *et al*. (2018). We tested for statistical differences in the ROC curves between different mapping designs using a Wilcoxon test on the distribution of simulated AUC values. We performed multiple-testing correction for the tests comparing AUC distributions using the Bonferroni method.

### Genomic inflation factor

We calculated the genomic inflation factor (GIF) to evaluate the role of mapping design and genetic architecture on the overall signal of association. We evaluated GIF on chromosomes linked and unlinked to causal loci. We calculated the GIF by correcting for sample size following Freedman et al (2004).

### Data availability statement

The raw sequence data for our six reconstructed real-life hybrid swarm individuals is available at SRA BioProject accession PRJNA691163. The most up-to-date code associated with this project is available on GitHub: https://github.com/cory-weller/HS-reconstruction-gwas and an archived version of our repository is available at Zenodo (DOI: 10.5281/zenodo.4472955). This repository includes a containerized version of the reconstruction pipeline along with a test data-set.

## Results

### Genome reconstruction for a Hybrid Swarm

First, we evaluated our most-likely ancestor selection algorithm, which selects a minimum representative set of Most-Likely-Ancestors (MLAs). This step required optimizing a discrimination threshold for likelihood values calculated with HARP (Kessner *et al*. 2013, see methods). We found that classifying any potential founder as an MLA by counting the number of chromosomal windows with observed likelihoods in the top 5% (threshold of 0.95) identified all potential founders with high accuracy (Supplemental Figure S7). The threshold of 0.99 performed slightly better when 128 founding lines were used to found a population, while the threshold of 0.95 performed slightly better with only 32 founding lines. HARP thresholds of 0.95 and 0.99 perform similarly well across a range of simulated population parameters, suggesting this is a reasonable starting point for optimizing the pipeline for other populations (Supplemental Figure S8A). Reducing sequencing coverage by a factor of ten (from 0.05X to 0.005X) resulted in similar levels of ancestor selection accuracy (Supplemental Figure S8B), demonstrating that additional coverage may not necessarily improve reconstruction accuracy. For simplicity, we next conducted all genome reconstructions with the 0.99 threshold at a simulated sequencing coverage of 0.05X.

Next, we evaluated the accuracy of genotype calls following genome reconstruction based on the most-likely ancestors identified for any individual chromosome. In general, genotype reconstruction from simulated F5 Hybrid Swarm individuals was very accurate: the median percent of sites with correctly estimated genotypes was greater than 99.9% whether the population was founded by 32 or 128 founding lines or made use of DGRP or simulated haplotypes (Figure 2, Supplemental Figure S10). For simulations founded by DGRP lines, 80.5% of reconstructed chromosomes from 32-founder populations exhibited > 99.9% accuracy, with the remaining 19.5% of reconstructions contributing to a long tail with a minimum of 84.5%. Increasing the number of founding lines to 128 resulted in genotype accuracy above 99% for all cases (minimum: 99.4%), with 83% of reconstructed chromosomes achieving greater than 99.9% accuracy. Reducing sequencing coverage by an order of magnitude from 0.05X to 0.005X resulted in more frequent over-estimation of recombination, though overall median genotype accuracy remained above 99% (Supplemental Figure S9). Using simulated haplotypes, we also show that reconstruction accuracy improves with genetic diversity (Supplemental Figure S10).

We also examined the accuracy of our estimates of recombination number, per individual. The number of recombination events estimated from was generally high but differed between Hybrid Swarm populations with different numbers of founders. Reconstructions of individuals from Hybrid Swarm populations with more founders generated more accurate estimates of recombination count (Lin’s concordance correlation coefficient: 98% and 50% for DGRP 128- and 32-way Hybrid Swarms, respectively; see Table 1). For 32-founder populations, DGRP-derived reconstructions tended to over-estimate recombination counts. Results on recombination count accuracy from simulated haplotypes are reported in Supplemental Table 1.

**Table 1.**
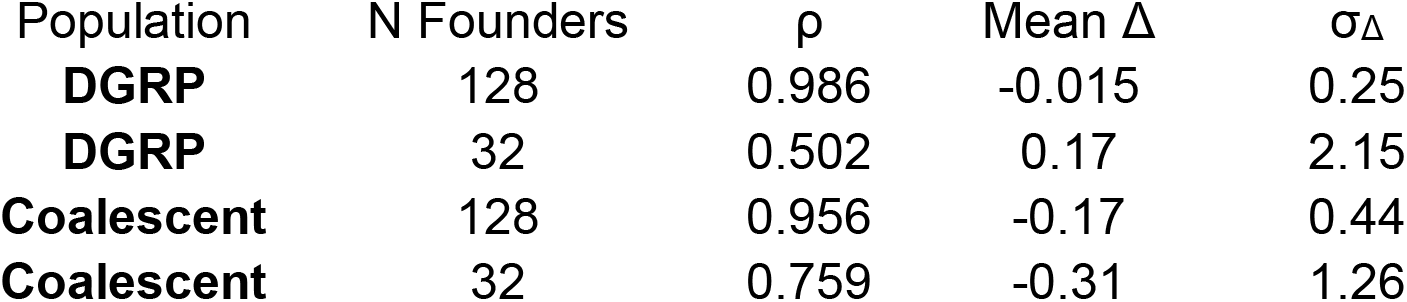
Accuracy of estimated number of recombination events following chromosome reconstruction from 0.05X sequencing data. A high concordance correlation coefficient (Lin’s ρ) indicates agreement between estimated and true recombination counts for 400 reconstructed chromosomes (coalescent-derived populations) or chromosome arms (DGRP-derived populations). Coalescent-derived populations are described across a range of values for effective population size N_e_ and mutation rate μ. Δ represents the difference between estimated and true recombination counts, and σ_Δ_ represents the mean of 400 standard deviations of Δ. Reconstructions were performed with a maximum of 16 most-likely-ancestors with a HARP threshold of 0.99 (see methods for more details). The coalescent-derived populations described here were simulated with an effective population size of N_e_=1×10^6^ and mutation rate μ=5×10^-9^.

| Population | N Founders | $\rho$ | Mean $\Delta$ | $\sigma_{\Delta}$ |
| --- | --- | --- | --- | --- |
| **DGRP** | 128 | 0.986 | -0.015 | 0.25 |
| **DGRP** | 32 | 0.502 | 0.17 | 2.15 |
| **Coalescent** | 128 | 0.956 | -0.17 | 0.44 |
| **Coalescent** | 32 | 0.759 | -0.31 | 1.26 |

Although genome reconstruction using RABBIT accurately calls genotypes, and generally estimates the number of recombination events well, we observe some reconstructions that are hyper-recombinant (i.e. > 10 recombination events for our simulations of an F5, Figures 2 and 3). These hyper-recombinant individuals had among the lowest genotyping accuracy, and this was the case for both DGRP-derived and coalescent-derived hybrid swarm individuals. The cause of these hyper-recombinant reconstructions is not always clear, but frequently appears due to RABBIT’s estimation switching between closely-related haplotypes. Because the number of inferred recombination events is determined by the number of generations of recombination, hyper-recombinant individuals (or regions of a genome) can be easily identified and removed as part of quality control (see Erickson et al 2020).

**Figure 3.**
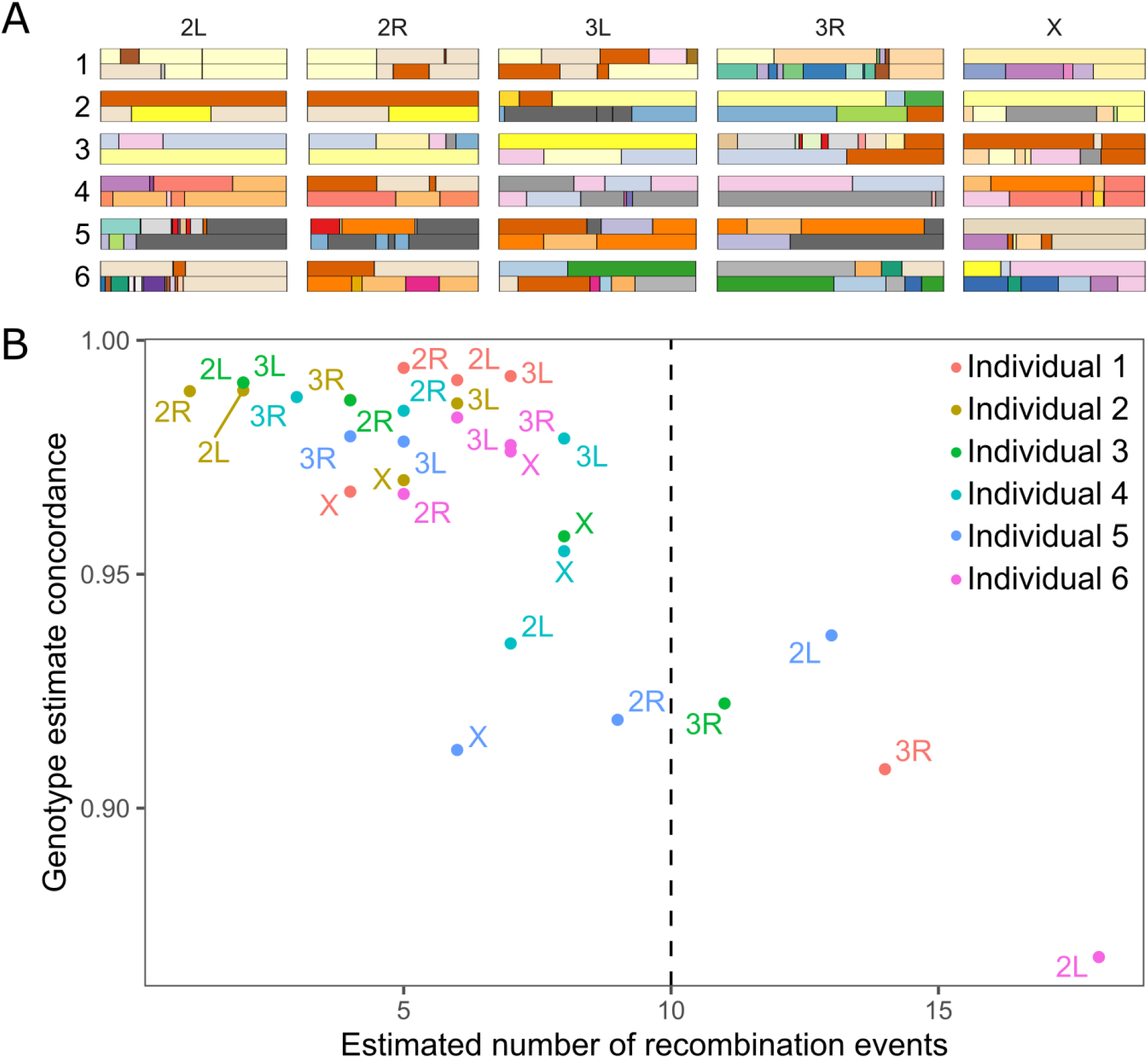
RABBIT chromosome reconstructions for real-life hybrid swarm individuals. A: Depictions of estimated haplotypes for six low-coverage F5 *Drosophila melanogaster*, identified by sample number. B: Genotype estimate concordance between our reconstruction pipeline and higher-coverage genotype calls. Colors correspond to individuals 1-6 in panel A. Our low-coverage genotype estimates tend to be 97-99% concordant with high-coverage calls, except in cases where RABBIT estimates are highly recombinant. A dashed line at 10 recombination events marks our threshold for excluding assumedly inaccurate hyper-recombinant reconstructions.

We contrasted the accuracy of genotype calls made with RABBIT to those made with an alternative tool, STITCH (Davies *et al*. 2016). STITCH does not rely on priors of parental haplotypes, but instead performs imputation and phasing based on patterns of linkage among the entire set of sequenced Hybrid Swarm recombinants. As expected, STITCH accuracy improved with greater sample size of *N* sequenced individuals (Figure S11), providing 68.5% genotype accuracy with *N*=100, increasingly approximately linearly until 94.5% accuracy at *N*=3000, and providing 99% accuracy with *N*=4000.

Genome reconstruction and genotype calls of real Hybrid Swarm individuals sequenced at ultra-shallow (~0.05X) and high (~8) coverage were highly concordant (~95%; Figure 3). Accuracy is particularly high for reconstructed chromosomes that predict fewer than 10 recombination events.

### GWAS accuracy

We evaluated GWAS accuracy for several configurations of Hybrid Swarms and compared these to simulations of GWAS using an 8-way RILs (modeled after the DSPR), and an inbred reference panel (the DGRP). Examples of representative Manhattan plots for a single causal locus of large effect are shown in Figure 4. RILs tend to result in a strong and wide association peak near the causal allele, while other populations result in only a small number of sites (typically on or neighboring the causal locus) with p-values below 10^-5^.

**Figure 4.**
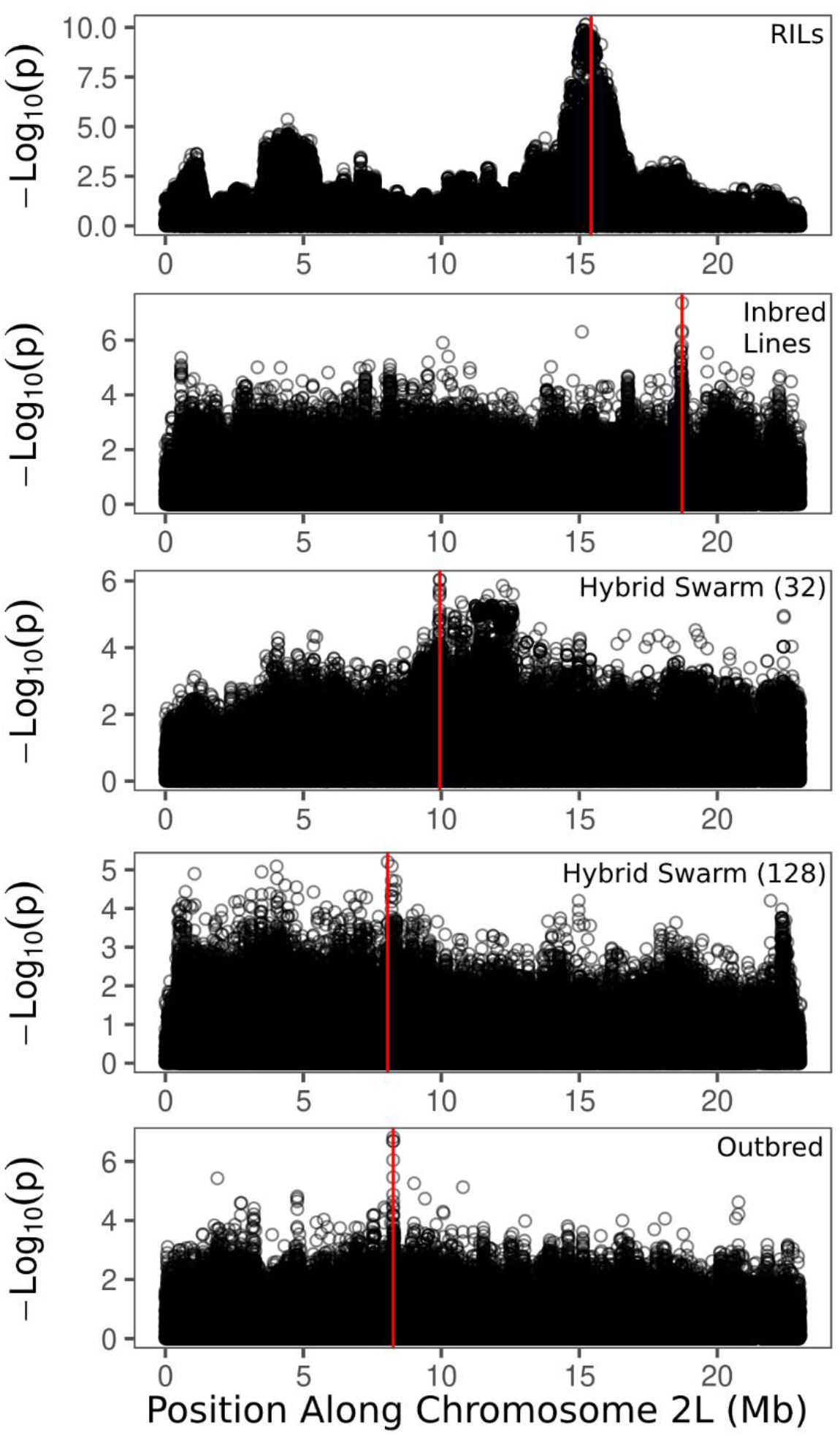
Examples of Manhattan plots from our simulated GWAS. These simulations correspond to a single causal locus of large effect, with position indicated by the vertical red line. Recombinant Inbred Lines tend to display a wide association peak near the causal locus, while other populations show only a small number of sites tightly clustered on the causal locus.

To assess overall performance of these mapping populations across hundreds of simulations, we generated Receiver Operator Characteristic (ROC) curves and calculated Area Under the Curve (AUC; Figure 5). In general, recombinant inbred lines outperformed other mapping designed. RILs were only significantly outperformed for a single-locus trait of large effect mapped in a 128-founder inbred reference panel, though both populations approach a median AUC of 1.0 (Figure 5B, median RIL AUC = 0.997, median 32-founder Inbred Line AUC = 0.999, p=1.101×10^-6^). Inbred reference panels consistently had the lowest AUC for multi-locus traits (Figure 5C-F). We suspect that the decreased accuracy of association mapping using inbred lines is due to linkage to causal sites, as well as spurious long-distance linkage (see below, *Genomic Inflation Factor*). See Supplemental Table S2 for comparisons between all mapping population types.

**Figure 5.**
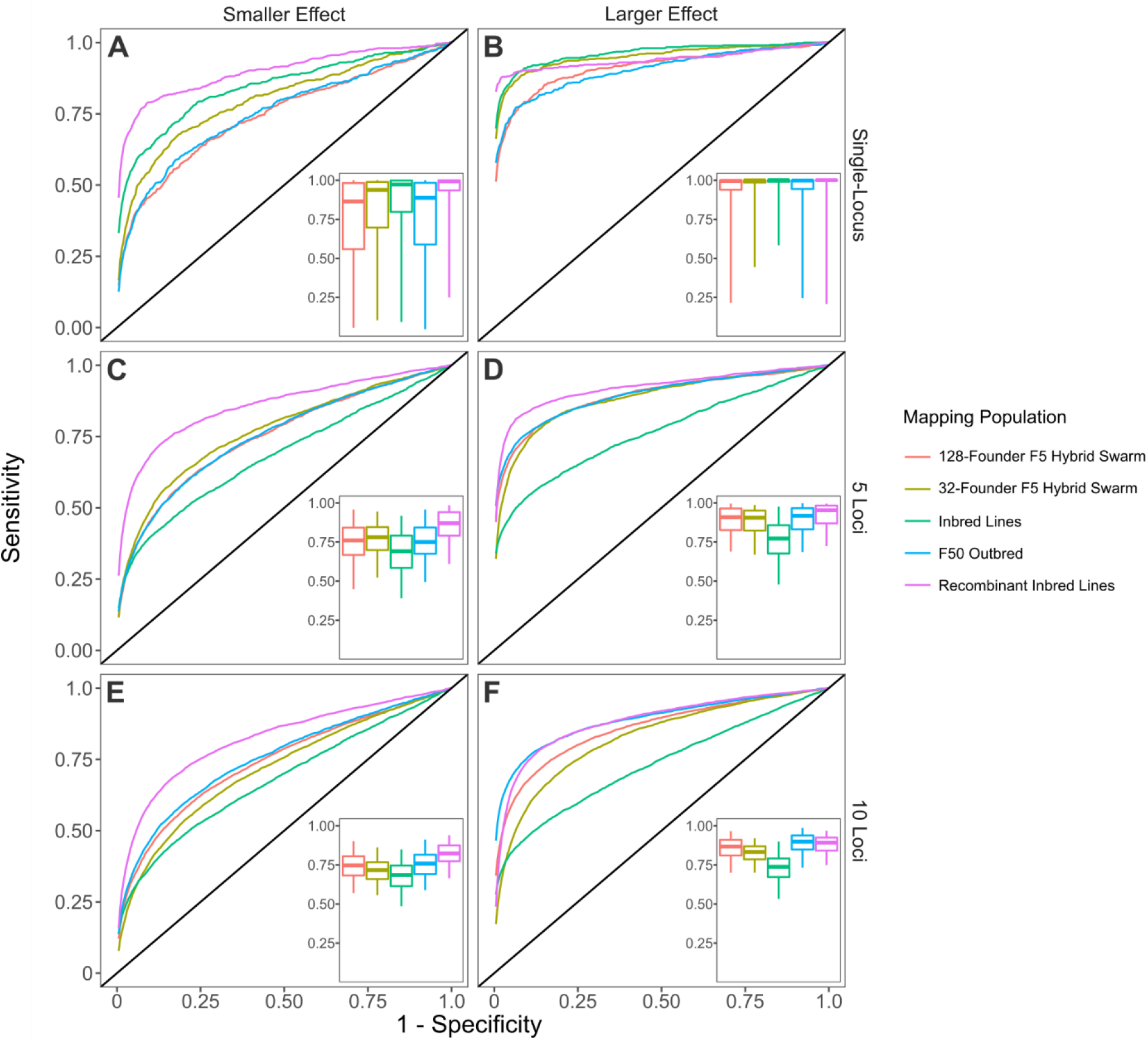
Receiver Operator Characteristic (ROC) curves for GWAS simulations. ROC curves represent the aggregate of 500 GWAS simulations (each comprised of 5000 individuals) per parameter combination. To generate single representative ROC curves for each population, we stepped through all specificity values and calculated the mean sensitivity. Inset boxplots display the distribution of AUCs, with whiskers spanning the middle 95% of data. See methods for descriptions for generating each mapping population.

Hybrid Swarm populations, composed of either 32- or 128-founders intermated for 5 generations had intermediate performance (Figure 5), and generally resembled fully outbred (F50) populations composed of 128 founders. We also evaluated the GWAS performance when crossing 32 or 128 founders to generate F1 and F2 mapping populations (Supplemental Figure S12). For single-locus simulations, the number of generations of recombination did not influence accuracy, although 32-founder populations performed better than 128-founders. For multi-locus simulations, GWAS accuracy improved with more generations of recombination, and this effect was more pronounced with 10-locus traits compared to 5-locus traits. Statistical tests comparing different models can be found in Supplemental Table S2.

### Genomic Inflation Factor

An alternative way to assess the quality of GWAS using different mapping designs is to compare the Genomic Inflation Factor. This metric describes how much the genome-wide distribution of test statistics differs from a null expectation. Values greater than 1 can indicate an excess of low p-values and can reflect population structure (Reich and Goldstein 2000) or polygenicity (Yang *et al*. 2011). We calculated the genomic inflation factor for any given mapping population and genetic architecture genome-wide (across all autosomes), on the autosome arm containing the causal allele (linked), and for sites on the autosome physically unlinked to the causal allele.

The relative ranking of GIF remained the same between a mono- and polygenic architecture with Inbred Lines producing the greatest GIF, and outbred F50 populations producing the lowest GIF (Figure 6). Hybrid Swarm populations showed intermediate GIF. Interestingly, Inbred Lines display elevated GIF even for sites physically unlinked to causal loci, suggesting spurious long-distance linkage disequilibrium. This inflation on unlinked chromosomes for inbred lines persists when filtering out sites below 5% frequency (Supplemental Figure S13), suggesting the pattern cannot be attributed to low-frequency alleles alone. Elevated GIF on unlinked chromosomes when using inbred panels such as the DGRP suggests caution in interpreting GWAS using this type of mapping population, and could contribute to the observed differences in mapping results of the same trait studied in both inbred lines and an advanced Hybrid Swarms derived from the DGRP (Huang *et al*. 2012b, 2020). The Hybrid Swarm, propagated for at least two generations, was able to decrease GIF at unlinked loci and by five generations the problem of elevated GIF on unliked chromosomes was ameliorated.

**Figure 6.**
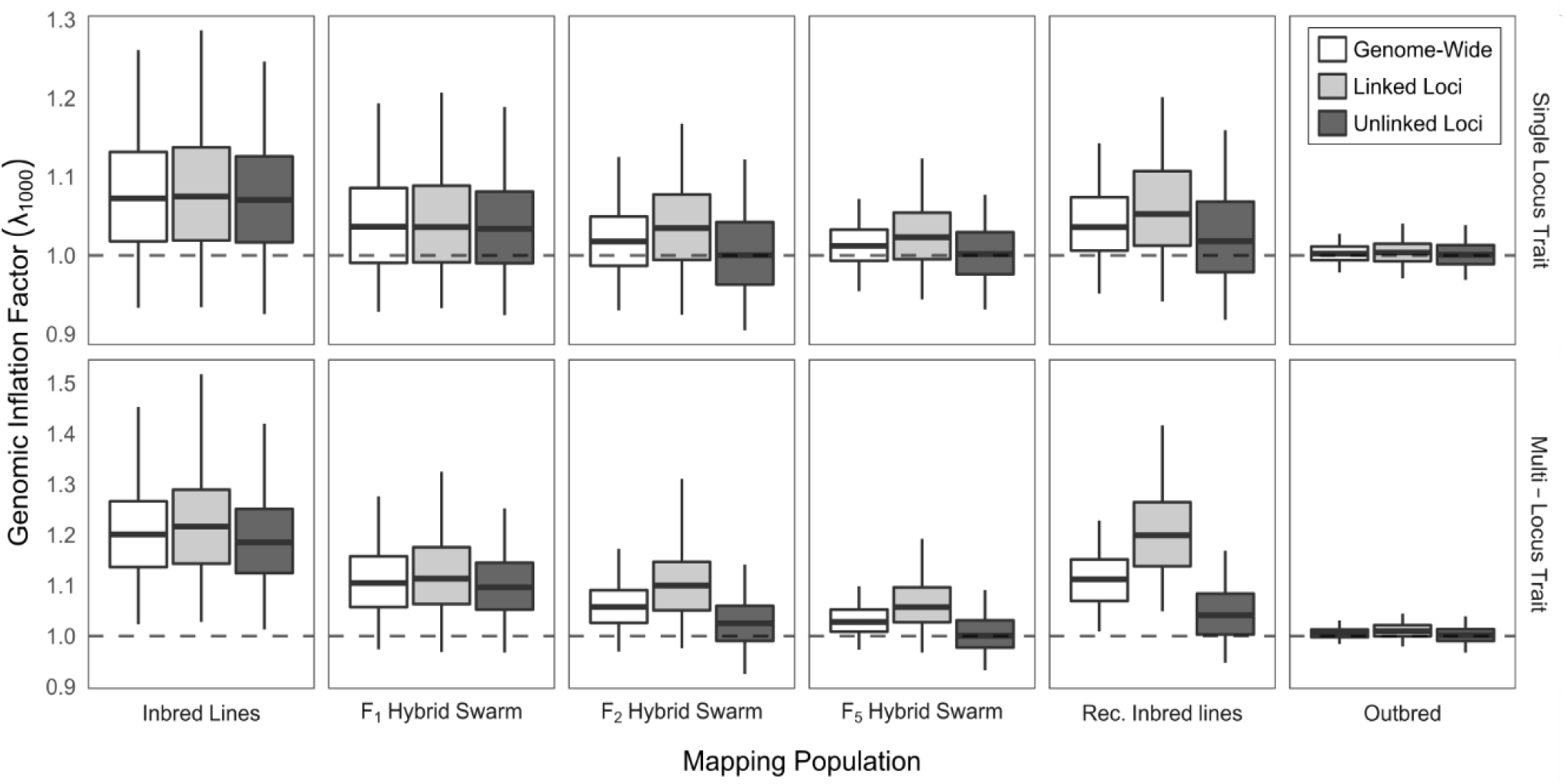
Genomic Inflation Factor (GIF, λ1000) for simulated GWAS with monogenic trait of with a large effect locus, or multi-locus trait of smaller individual effects. GIF is calculated genome-wide (across all autosomes); on the autosome arm containing the causal allele (linked); and for sites on the autosome physically unlinked to the causal allele. λ is calculated as the ratio of observed to expected χ2 values, and a correction is performed to produce the null expectation of a sample size equal to 1000 individuals (see Materials and Methods for details). Data are averaged over 500 GWAS simulations (each comprised of 5000 individuals), with phenotypes assigned in a casecontrol framework. Boxes represent the median and interquartile range; whiskers extending to the lower and upper bounds of the 95% quantiles. Hybrid Swarm populations here used 128 founders. Outbred: F50 population founded by 128 inbred lines.

## Discussion

Here, we show that the Hybrid Swarm is a viable method for association mapping using experimental outbred populations. We first demonstrate that ultra-low coverage, individual-based whole genome sequencing can be used to reconstruct accurate diploid genomes for species across a range of diversity levels (Figure 2, Supplemental Figure S10), and we validate the quality of genome reconstruction using real Hybrid Swarm individuals sequenced at both high and low coverages (Figure 3). Genome reconstruction with a reduced subset of most-likely-ancestors is computationally efficient (Supplemental Figure S2), and not dependent upon the number of individuals genotyped unlike STITCH (Supplemental Figure S11), enabling a wide range of experimental designs. Next, we demonstrate that association mapping using the Hybrid Swarm can match or outperform various types of inbred lines and completely outbred populations (Figures 4, 5, Supplemental Figure S12, Supplemental Table S2). Lastly, we have described a set of computational tools to simulate various mapping populations (including genome reconstructions, and GWAS), enabling this method to easily be applied to a variety of organisms.

### Considerations of the Hybrid Swarm approach

The Hybrid Swarm approach is applicable to a wide variety of organisms and experimental designs, conferring potential benefits over inbred reference panels. These benefits are realized in three primary ways by: 1) allowing researchers to address questions that require heterozygotes; 2) reducing labor and the influence of block-effects; and 3) breaking down linkage-disequilibrium and population structure that arises from the sampling of individuals from the wild. These benefits are possible due to the ability to accurately reconstruct genomes from ultra-low-coverage sequencing data, i.e. from cost-effective diluted DNA library preparation for a large number of individuals (Baym *et al*. 2015).

We evaluated simulated Hybrid Swarms founded by 32 or 128 founders to determine the effect of founding haplotype count on reconstruction accuracy and GWAS performance. One might expect 32-founder populations to yield more accurate reconstructions than 128-founder populations, because it is simpler to make a correct inference out of a smaller pool of ancestors. Yet, 32-founder populations were less accurate in our DGRP simulations (Figure 2), and for some coalescent-based simulations (Supplemental Figure S10). The difference in reconstruction accuracy is perhaps due to the inverse relationship between SNP frequency and information content. In a 128-way cross, rare information-rich SNPs are only likely to exist and be sampled during low-coverage sequencing if nucleotide diversity is high. The DGRP haplotypes happen to exist in a parameter space with higher nucleotide diversity, resulting in 128-way reconstructions being more accurate. For GWAS, 32-founder Hybrid Swarm populations tended to perform better than 128-founder populations if the simulated trait was controlled by a single locus (Figure 5). Taking these two factors into consideration, broadly speaking, a Hybrid Swarm with fewer founding haplotypes may be more appropriate for reconstructing genomes when genetic diversity in the founding lines is low, or if the trait of interest is presumably controlled by one or few loci. Conversely, a greater number of founders may be desired when genetic diversity is high, or the trait of interest is likely to be complex.

The Hybrid Swarm method is not limited to populations founded by inbred lines, as the technique can be applied to populations where phased genomes are available for all outbred founders. Phasing the outbred founders could be accomplished in two basic ways. For instance, phased genomes could be generated using a variety of long-read sequencing technologies (Pollard *et al*. 2018). Alternatively, trio- and quartet-phasing methods (Patterson *et al*. 2015), coupled with high coverage sequence data from the outbred parents plus a limited (≤2) number of F1 offspring, would enable accurate phasing of parents to be used for reconstruction of downstream recombinant genomes.

One experimental consideration is the effects of drift and selection that operates during the generation of the Hybrid Swarm (e.g., Thépot *et al*. (2014)). However, it is not clear that such processes will affect the ability to reconstruct genomes nor the specific outcome of GWAS, unless the trait of interest was under strong selection itself. What may be more likely is the reduction of haplotypic diversity due to drift or selection will slightly decrease power to detect associations, as we see in the differences between 32- and 128-way crosses (Supplemental Figure S10). The distribution of haplotypes can also be skewed by line-specific differences in fitness or fecundity, with such differences being observed for DGRP lines (Horváth and Kalinka 2016). To attenuate haplotype dropout, it may be prudent to seed a Hybrid Swarm with a large population of directed F1 hybrids produced by round-robin crosses. The F1 population would then be followed by a limited number of generations (e.g., 4-5) of random outbreeding. This approach was used by Erickson *et al* (2020) and resulted in a relatively even distribution of founding haplotypes after 5 generations.

We compared genotype calls from our low-coverage reconstruction pipeline to genotype calls from higher coverage data for six experimental Hybrid Swarm individuals (Figure 3). Our results show high concordance between reconstructed genomes and high coverage genotypes, particularly if reconstructions with 10 or more inferred recombination events are excluded. Researchers wishing to minimize the impact of inaccurate genotype estimates could implement a stringent limit on inferred recombination counts.

### Representation of heterozygotes

One clear difference between inbred and outbred mapping populations is the presence of heterozygotes. On the one hand, the presence of heterozygotes in outbred populations decreases power to detect association relative to inbred lines for an (semi-) additive allele with a given effect size (Figure 5). This basic statistical effect may also be influenced by differences in realized genetic variance between inbred and outbred populations (Genissel *et al*. 2004). However, the reduced statistical power of association mapping in outbred populations may be ameliorated by reduced inbreeding depression (Lee *et al*. 2017) and by the ability to assess the heterozygous effects of alleles.

The ability to assess heterozygous effects of alleles will provide valuable insights into several interesting aspects of biology, such as the nature of dominance and the identity of regulatory polymorphisms governing allele-specific expression in a variety of heterogenous genomes. An increased understanding of dominance relationships and regulatory polymorphisms is important for advancing our understanding of quantitative trait variation and evolution. For instance, several theoretical models have shown that context dependent dominance of quantitative fitness traits can underlie the stable maintenance of polymorphisms subject to seasonally variable (Wittmann *et al*. 2017) or sexually antagonistic (Connallon and Chenoweth 2019) selection. The ability to efficiently map loci with context dependent dominance relationships will aid in the understanding of the stability and abundance of polymorphisms maintained by these forms of balancing selection.

Regulatory polymorphisms are known to underlie genetic variation in expression (Brem *et al*. 2002; Cavet *et al*. 2003; Rockman and Kruglyak 2006) and this expression variation can potentially be resolved to exact nucleotide differences (Grosveld *et al*. 1987; Rave-Harel *et al*. 1997; Bosma *et al*. 2002). The resulting differences in expression can manifest as phenotypic changes to drive local adaptation (Kudaravalli *et al*. 2009; Fraser *et al*. 2010; Fraser 2011, 2013). Allele-specific expression (ASE) arising from cis-acting regulatory factors is a common mechanism to produce heritable differences in expression (Yan *et al*. 2002; Cowles *et al*. 2002; Lo *et al*. 2003; Doss 2005). Because allelic expression biases are only produced (and detectable) in heterozygotes, Hybrid Swarm populations facilitate the study of regulatory genetic variation (i.e. ASE) as a driver of local adaptation in a variety of organisms.

### Undirected outbreeding in a common environment

The Hybrid Swarm approach involves propagation of a single large outbred population via undirected crossing. This design confers benefits over alternatives of either rearing inbred lines separately or performing controlled crosses. The relative value of these benefits may vary across organisms and experimental designs.

Individuals within a Hybrid Swarm are reared in a common environment. This reduces the influence of random block effects associated with rearing families or closely related individuals in separate enclosures or defined areas. Therefore, studies examining genotype-by-environment interactions can dramatically expand the number of environmental factors used because all individuals were previously reared in a common environment, enabling a Hybrid Swarm to be randomly scatted across environmental treatments. This feature is in contrast to rearing each inbred family across all environments, a practice which generally limits the number of environmental treatments used. Rearing individuals in a common environment also likely reduces genotype-environment correlation that could be exacerbated by vial effects.

Random outbreeding of a single population can require less labor compared to performing controlled crosses or serial propagation of inbred lines. The relative value of these benefits is likely to be species specific. For instance, for larger animals (e.g., mice), controlled crosses may be logistically easier than a randomly mating swarm. For many other species, however, the Hybrid Swarm would be logistically advantageous. The cost of maintaining inbred reference panels also varies among taxa, most notably between those species which can be kept as seeds or in a cryogenically preserved state. For other species, such as flies, the monetary and environmental cost of maintaining hundreds to thousands of inbred lines may be prohibitive.

One drawback draw-back of the Hybrid Swarm is the lack of genetic replicates for phenotyping. Inbred panels allow for a single genotype to be phenotyped multiple times, reducing the effects of error associated with phenotyping and micro-environmental variation (Mackay and Huang 2018). Because this benefit is typically not possible in the Hybrid Swarm (unless clones can be propagated), the Hybrid Swarm method represents a tradeoff, reducing the influence of block effects while increasing error associated with phenotyping.

### Hybrid Swarm breaks down population structure and linkage disequilibrium

Recombination between lines in the Hybrid Swarm approach allows for greater dissection of functional polymorphisms segregating between genetically structured populations. If an association study incorporates haplotypes from multiple distinct source populations, causal variants would segregate along with other linked variants. Thus, to identify genetic mechanisms of local adaptation and trait variation in general, it is necessary to minimize false positives from linked non-causal loci. Corrections due to relatedness can reduce the type I error rate to some degree (Yu *et al*. 2006; Price *et al*. 2010; Yang *et al*. 2014), and can be further reduced by a greater extent of recombination. Therefore, the Hybrid Swarm approach allows researchers to tailor the mapping population to the traits of interest, by perhaps selecting founders from diverse origins to test specific hypotheses about the distribution of functional variation in the wild.

A limited number of rounds of recombination during the propagation of the Hybrid Swarm also reduces long-distance linkage disequilibrium within a panel of inbred lines derived from a single locality. Long-distance LD results from correlated occurrence of rare variants (Huang *et al*. 2014), potentially contributing to false positives in GWAS (Figure 6). Our simulations showed a genome-wide inflation of p-values, even across physically unlinked chromosomes, for inbred panels and this pattern persisted even when rare variants were excluded (Supplemental Figure S13). Two or more generations of recombination were sufficient to reduce this inflation (Figure 6), at least when using association statistics that do not explicitly account for genetic structure in the sample. Whether such long-distance linkage disequilibrium substantially affects the results of GWAS in practice, remains an open question.

### Conclusions

Here, we demonstrate the feasibility of genome-reconstruction in Hybrid Swarm populations derived from many founding haplotypes and evaluate the power of this approach for association mapping. Our results suggest that the Hybrid Swarm approach can be a useful alternative to other MPP breeding designs and is applicable to model and non-model organisms.

## Author Contributions

**CW**: conceptualization, data curation, methodology, formal analysis, software, validation, visualization, writing the manuscript; **ST**: investigation, review of manuscript; **SR**: investigation; **AOB**: conceptualization, funding acquisition, project administration, resources, supervision, review and editing of the manuscript.

---

**Supplemental Figure S1.**
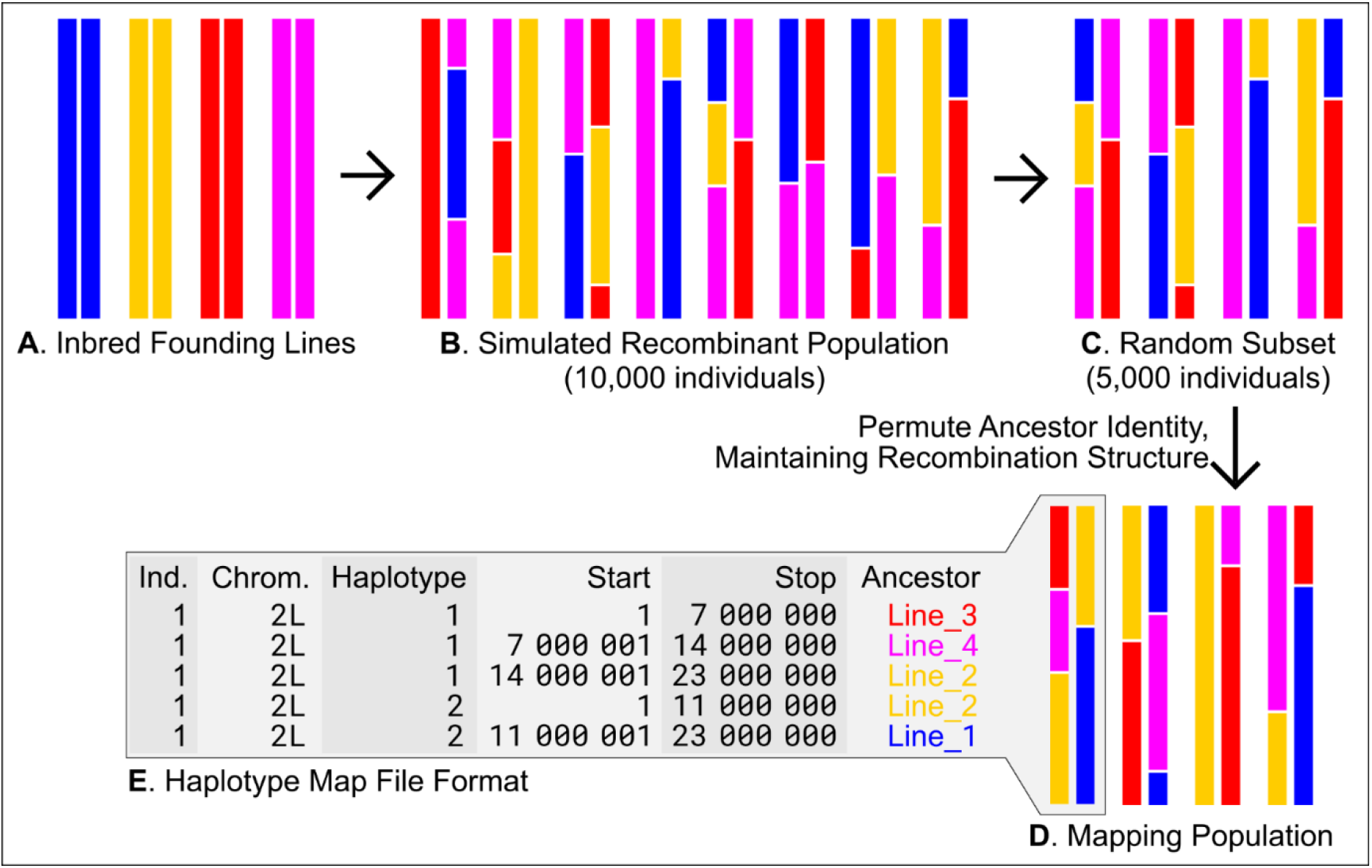
Basic structure of the forward simulator pipeline. Inbred founding lines (**A**) are randomly intercrossed to produce a recombinant population (**B**). Rapid generation of independent mapping populations is achieved by random down-sampling (**C**) and permutation of ancestry (**D**). Population genetic data is encoded in a highly compressed format **(E)** that references the positions of haplotype blocks instead of genotypes at every site, enabling us to generate 500 mapping populations for a given parameter combination.

**Supplemental Figure S2.**
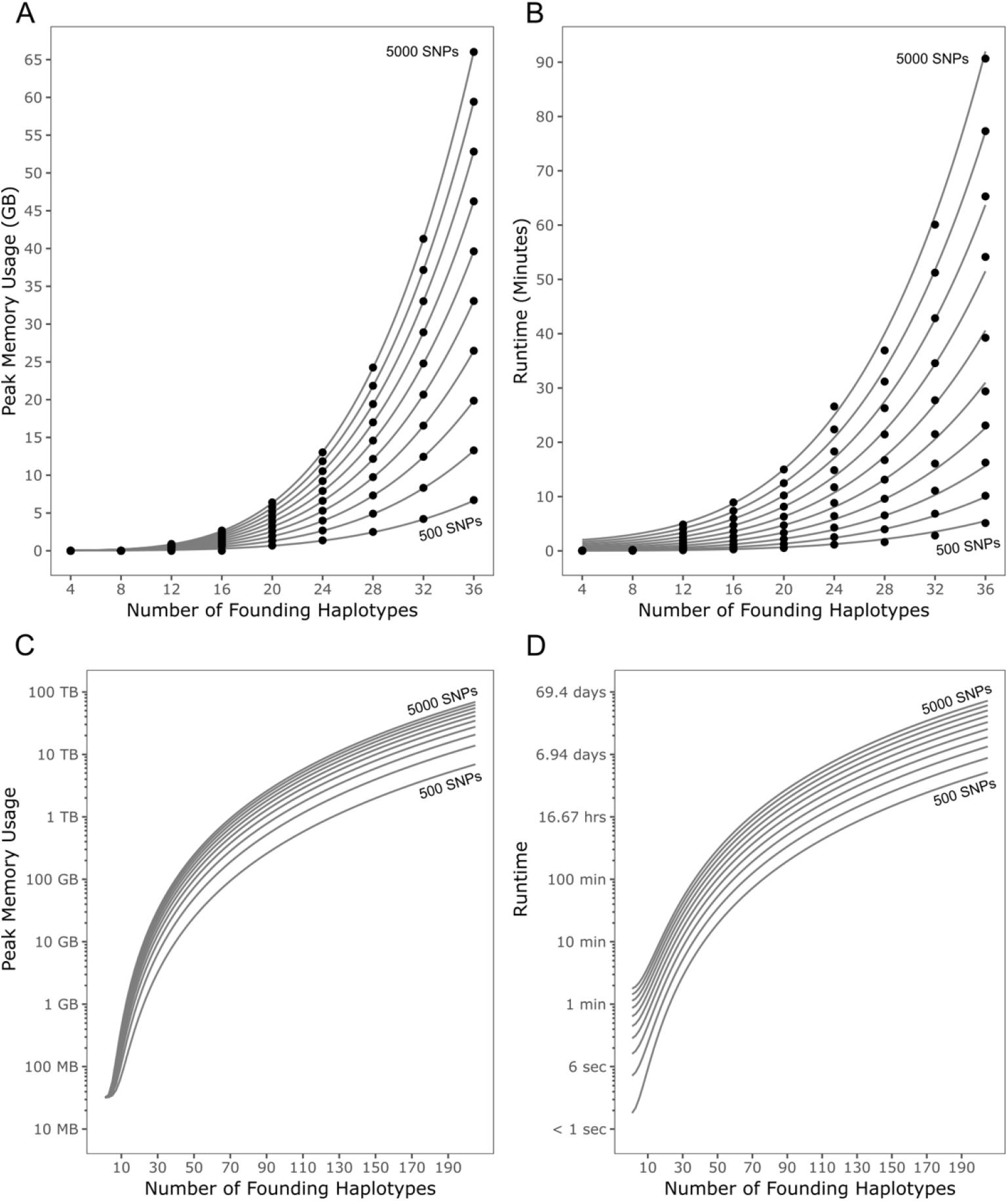
Resource usage of RABBIT during haplotype reconstruction. All reconstructions involve the same simulated 2L chromosome arm comprised of four haplotypes. Simulations included varied numbers of founding haplotypes (*N*) and a randomly selected set of markers (number of SNPs, *S*, incremented in steps of 500). All simulations included, at minimum, the four true haplotypes for the simulated individual. In **A** and **B**, points depict the mean of empirical values (over 10 replicates) and gray lines depict the defined regression models. Regression models for memory usage and runtime are extrapolated over a greater range for number of founding haplotypes in **C** and **D**, respectively. Peak memory grew linearly with number of SNPs used, and at a greater-than-linear rate with haplotypes (A). The runtime of RABBIT increased at a greater-than-linear rate for both number of SNPs and number of haplotypes, with the N parameter contributing most to resource cost (B). These models allowed us to estimate resource requirements at greater numbers of haplotypes (C & D) which would be unfeasible to measure empirically.

**Supplemental Figure S3.**
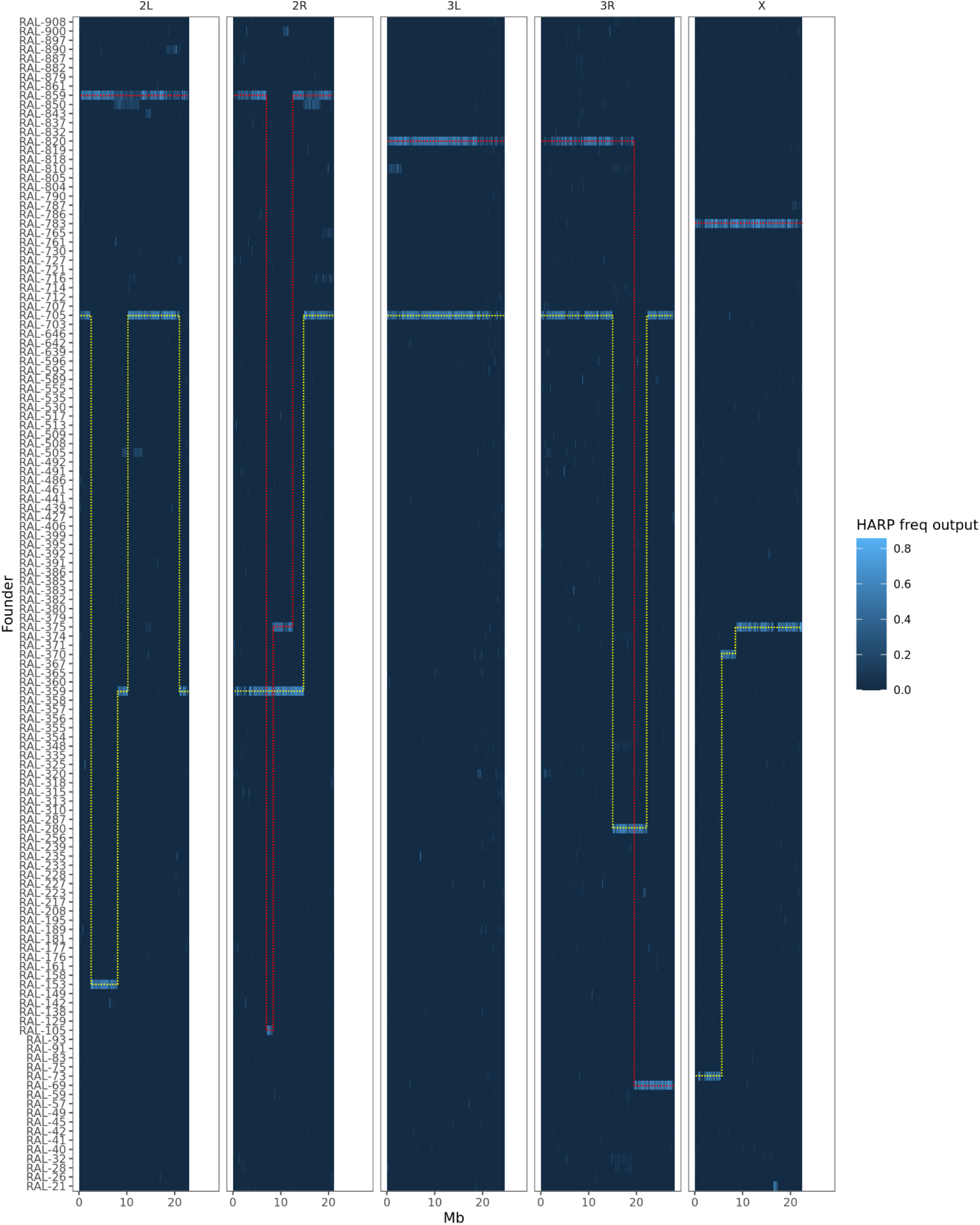
Example of unambiguous haplotype inference from HARP. For each chromosome, each haplotype is indicated by red and yellow dotted lines.

**Supplemental Figure S4.**
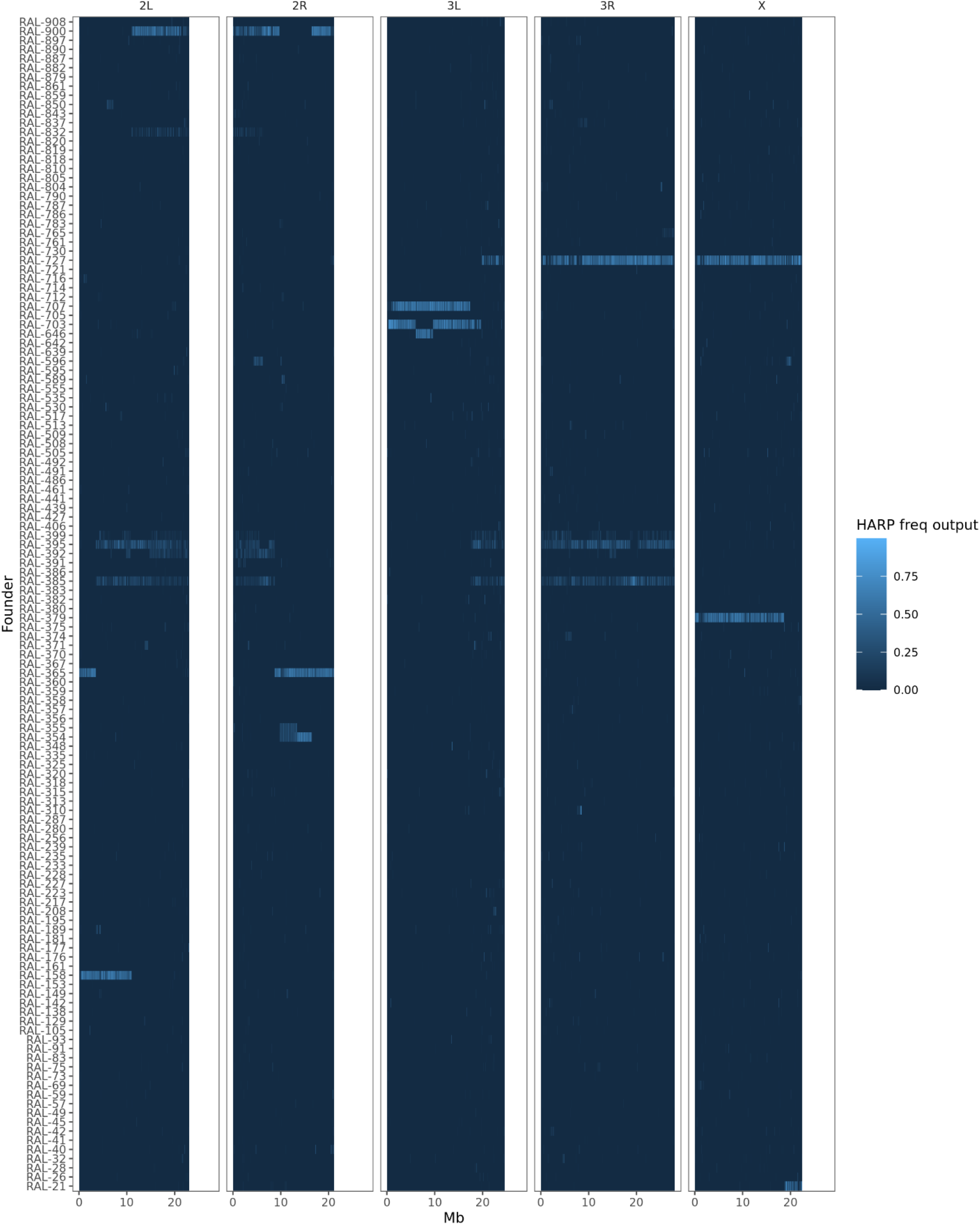
Example of HARP freq output with ambiguous haplotype inference. For each chromosome, each haplotype is indicated by red and yellow dotted lines. For this individual’s autosomes, at least one haplotype cannot be unambiguously resolved.

**Supplemental Figure S5.**
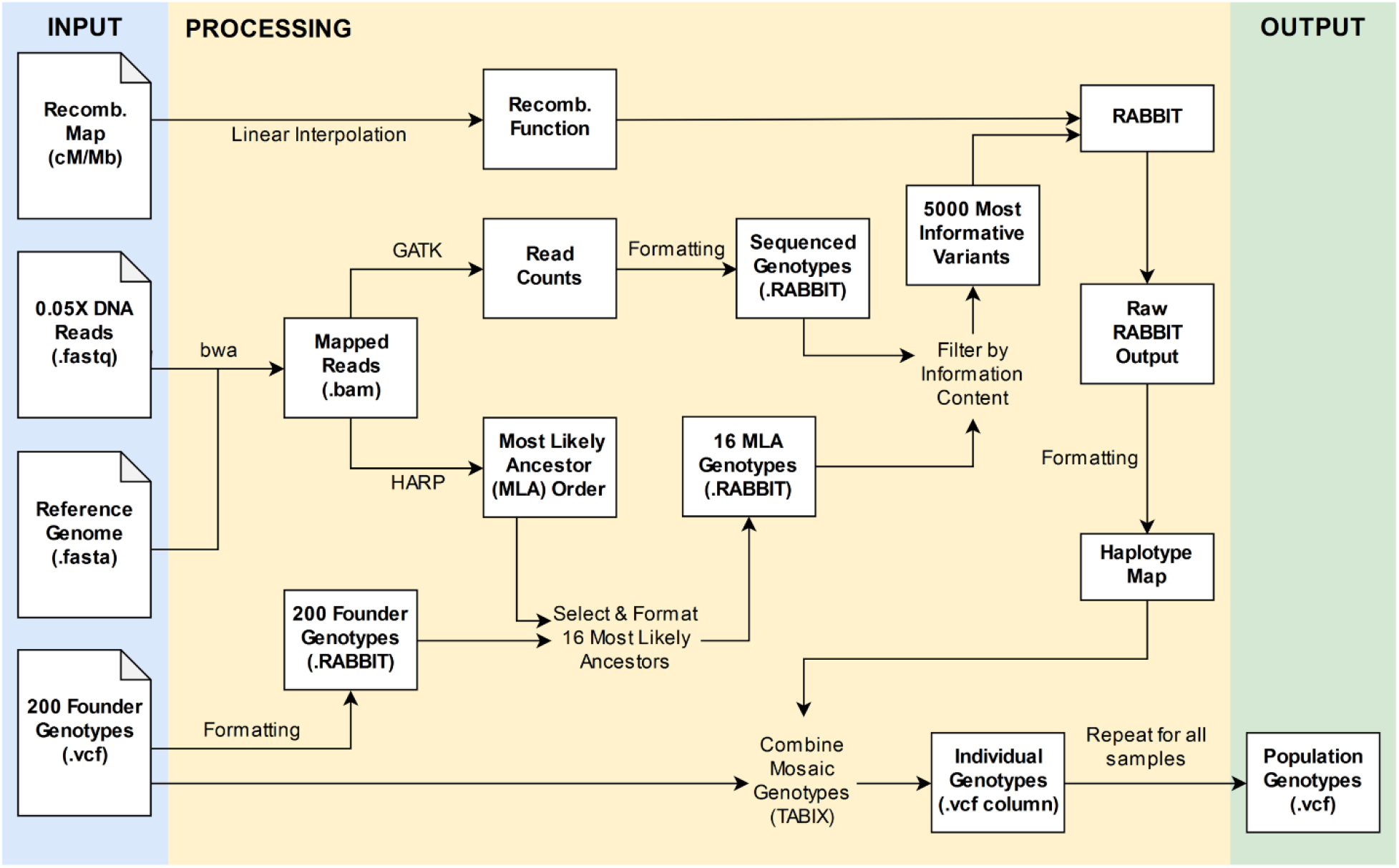
Workflow for reconstructing whole genomes from ultra-low coverage sequencing data. Three required inputs are sequencing reads (.fastq), a reference genome (.fasta), and homozygous (or phased) founder haplotypes (.vcf file). A recombination map can optionally be provided for use with RABBIT to inform variation in recombination rates across chromosomes. Flowchart designed in draw.io.

**Supplemental Figure S6.**
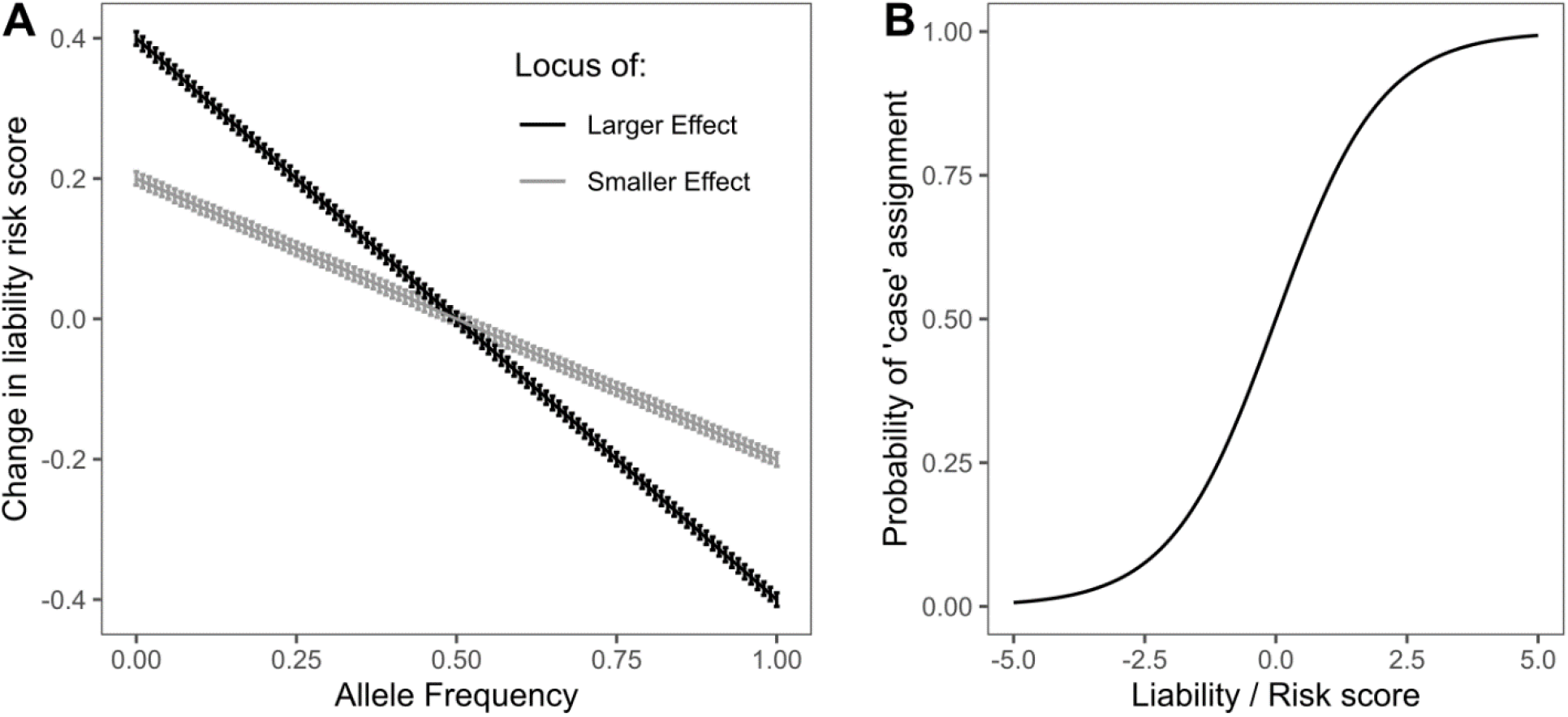
Liability phenotype model for GWAS simulations. for transforming additive risk scores to probability of assignment to the case population, i.e. probability of developing a disease. Rare alleles increase risk score, which in turns increases probability of being assigned a case phenotype. **A**: The relationship between allele frequency and effect on liability risk score. Simulations were modeled as larger or smaller effect with a user-defined effect size coefficient. Error bars describe the standard deviation of a gaussian sample which constitutes random phenotypic noise. **B**: Logistic transformation of the form y=(1+*e*^-x^)^-1^ translates the liability score into probability of being assigned to case population.

**Supplemental Figure S7.**
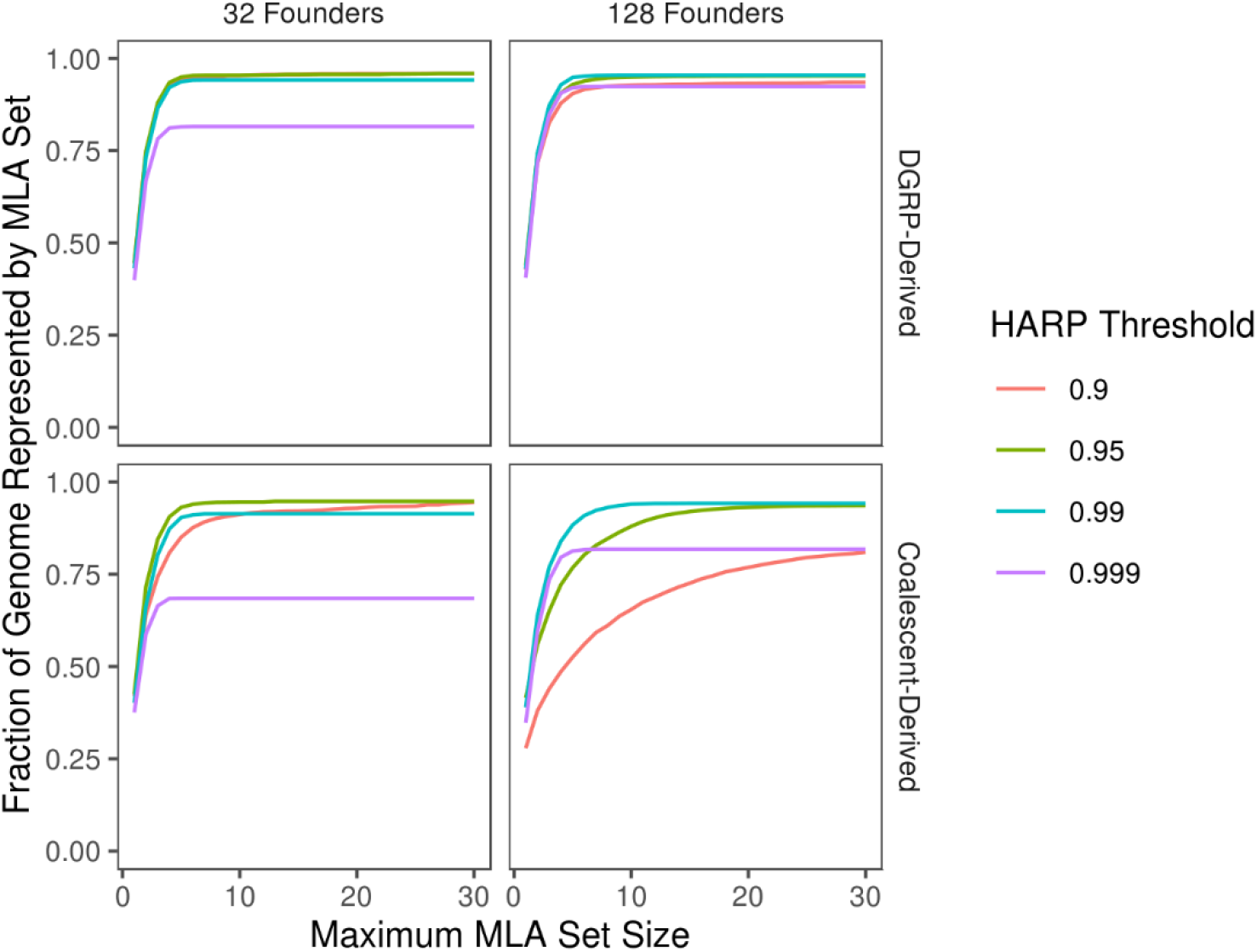
Optimization curves for Most-Likely-Ancestor (MLA) selection. To ensure reconstruction was both accurate and computationally feasible, we selected the smallest set of most-likely-ancestors that still represent the greatest fraction of to-be-reconstructed genomes. Increasing the upper limit for the number of MLAs chosen improves the fraction of genome represented with diminishing returns. The HARP threshold of 0.99 performed best for 128-founder populations, for both DGRP-derived and coalescent-derived simulations. Conversely, 32-founder populations performed best with a HARP threshold of 0.95. Data shown reports means across 400 replicates made up of 100 simulated individuals (4 autosomes each for coalescent simulations, 4 autosome arms each for DGRP simulations) per parameter combination. Coalescent-derived populations described here wer1e simulated with N_e_=1×10^6^ and mutation rate μ=5×10^-9^.

**Supplemental Figure S8.**
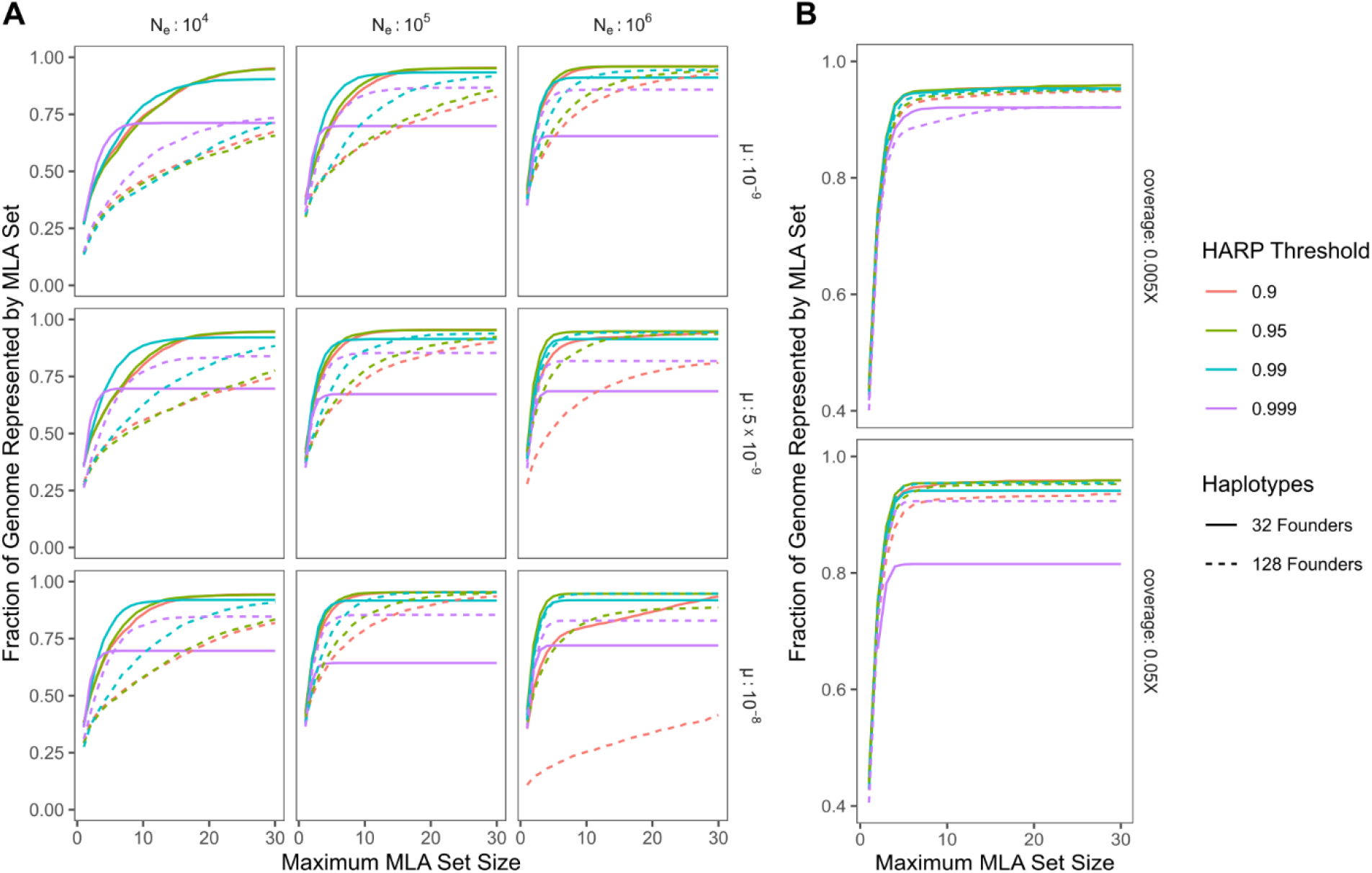
Optimization curves for Most-Likely-Ancestor inclusion for additional parameter combinations. A: SCRM (coalescent) derived populations across different parameters; B: accuracy is similar for both 0.05X and 0.005X sequencing coverage in populations simulated with DGRP genomes.

**Supplemental Figure S9.**
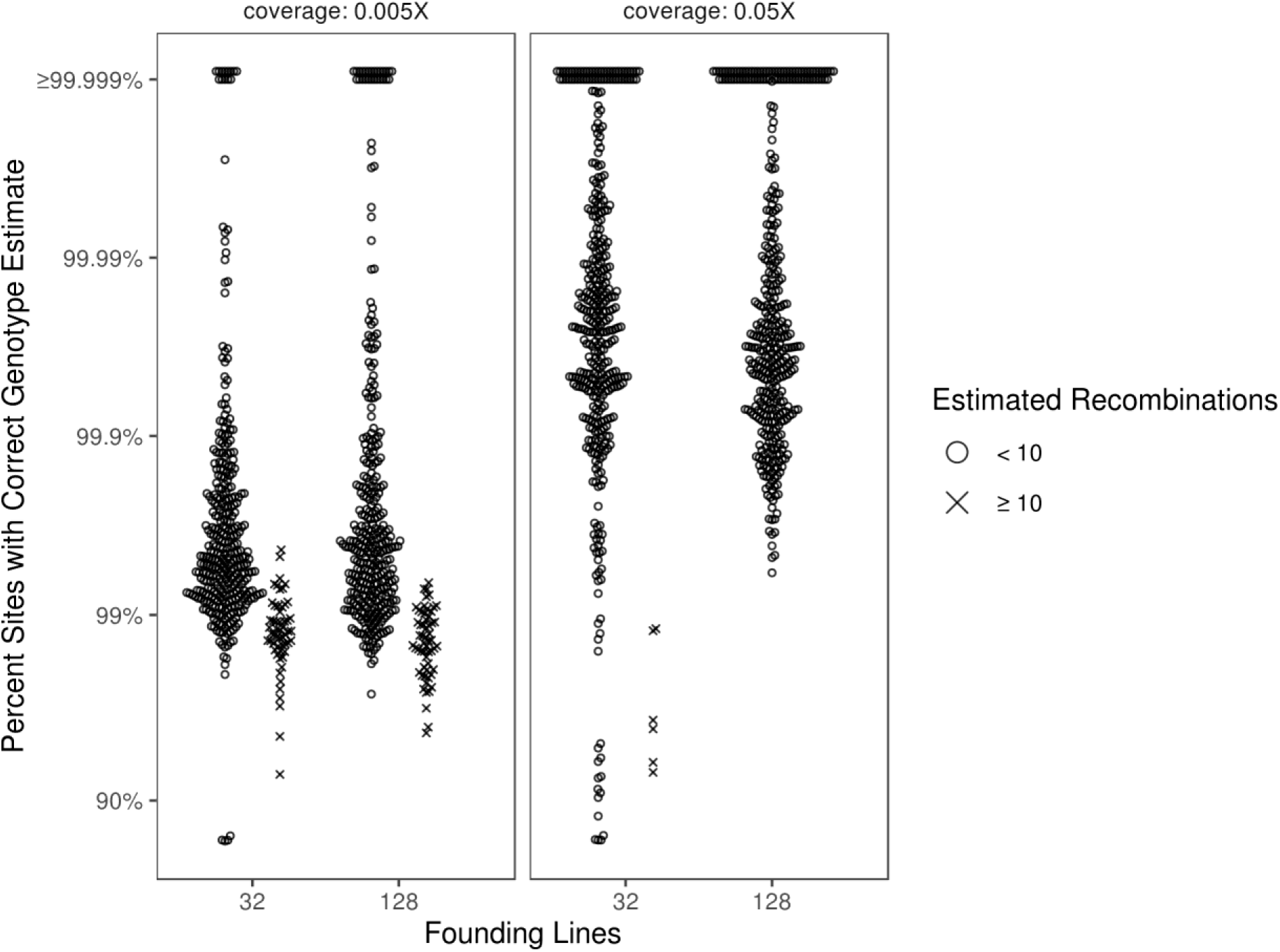
Accuracy of genome reconstruction for simulated, DGRP-derived F5 hybrid swarm individuals. Reconstructions were performed for populations simulated as being founded by either 32 or 128 inbred lines for two levels of ultra-low sequencing coverage. Accuracy is represented on a logit scale, as most points occur above 90%. Reconstructed chromosomes estimated to have ≥10 recombination events are denoted by an ×, offset from the bulk of the distribution. Each parameter combination includes 400 reconstructed chromosomes (from 100 simulated individuals).

**Supplemental Figure S10.**
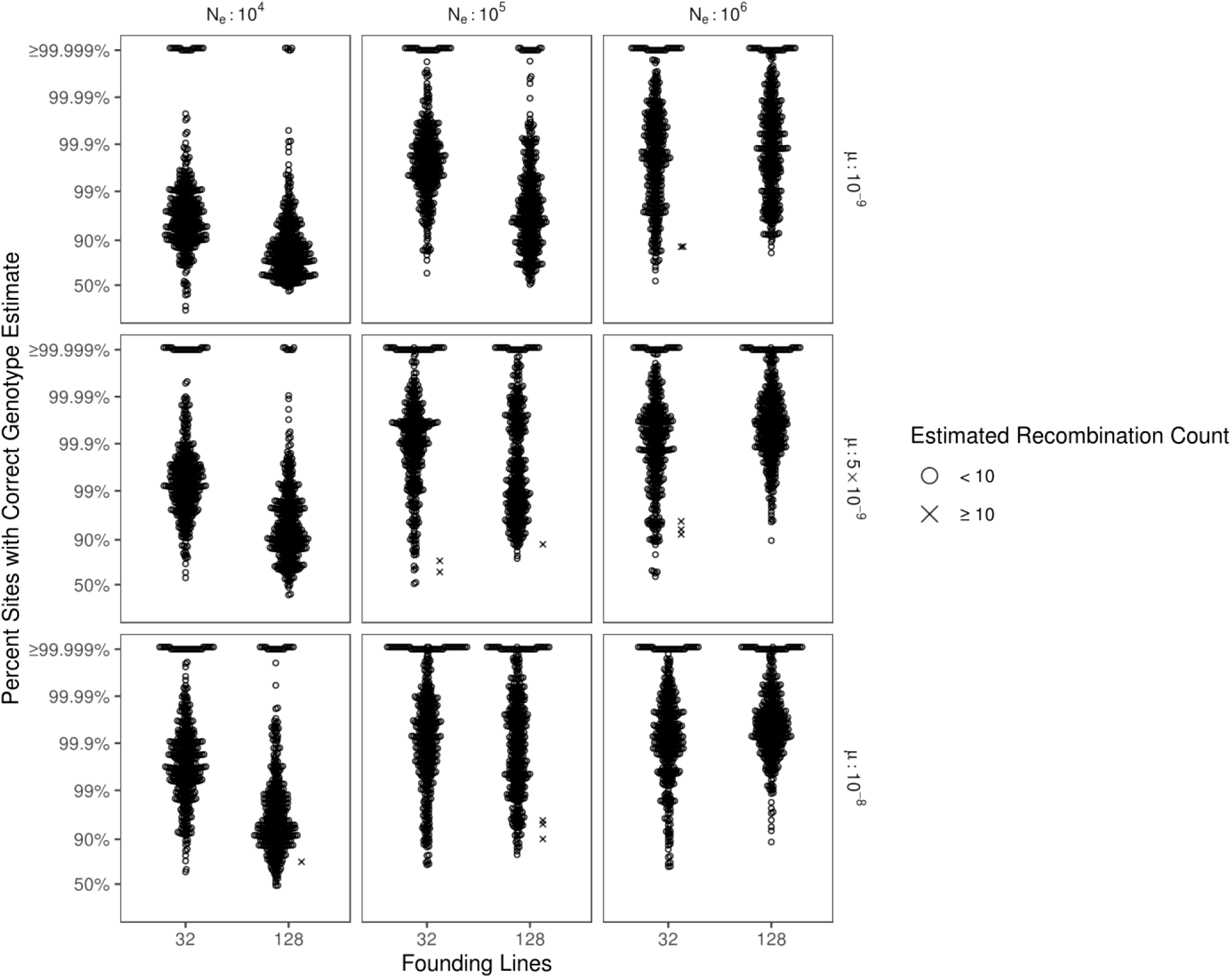
Accuracy of genome reconstruction for simulated, coalescent-derived F5 hybrid swarm individuals. Reconstructions were performed for populations simulated as being founded by either 32 or 128 inbred lines for various effective population sizes (Ne) and mutation rates (μ). Accuracy is represented on a logit scale, as most points occur above 90%. Reconstructed chromosomes estimated to have ≥10 recombination events are denoted by an ×, offset from the bulk of the distributions. Each parameter combination includes 400 reconstructed chromosomes (from 100 simulated individuals).

**Supplemental Figure S11.**
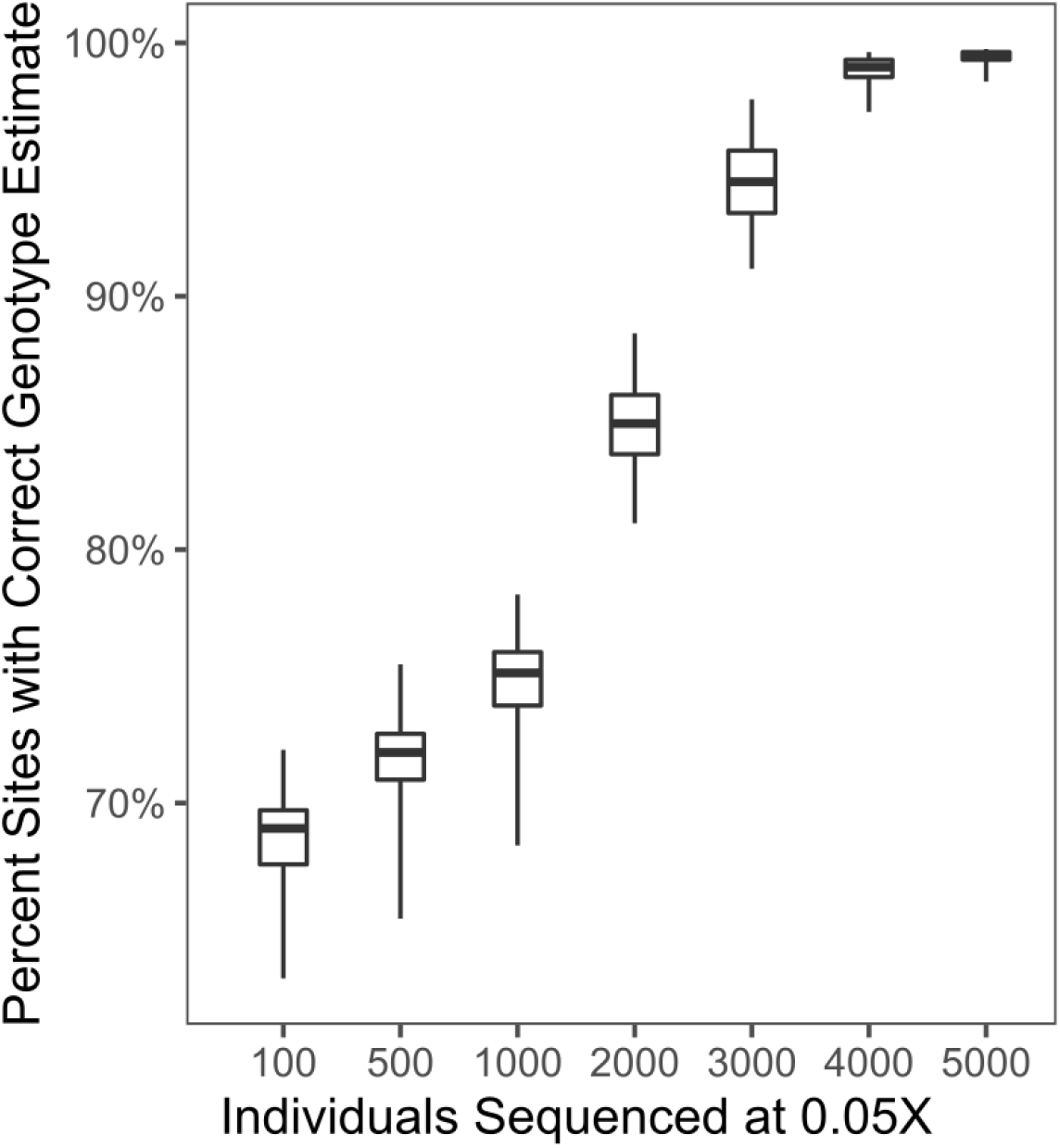
Genotyping accuracy of STITCH for simulated 32-founder F5 hybrid swarm populations sequenced at 0.05X coverage. Genotype accuracy improves with greater numbers of sequenced individuals, as STITCH infers missing genotypes from other haplotypes in the population. Boxes represent the median and interquartile range; whiskers extending to the lower and upper bounds of the 95% quantiles. See methods for parameters used.

**Supplemental Figure S12.**
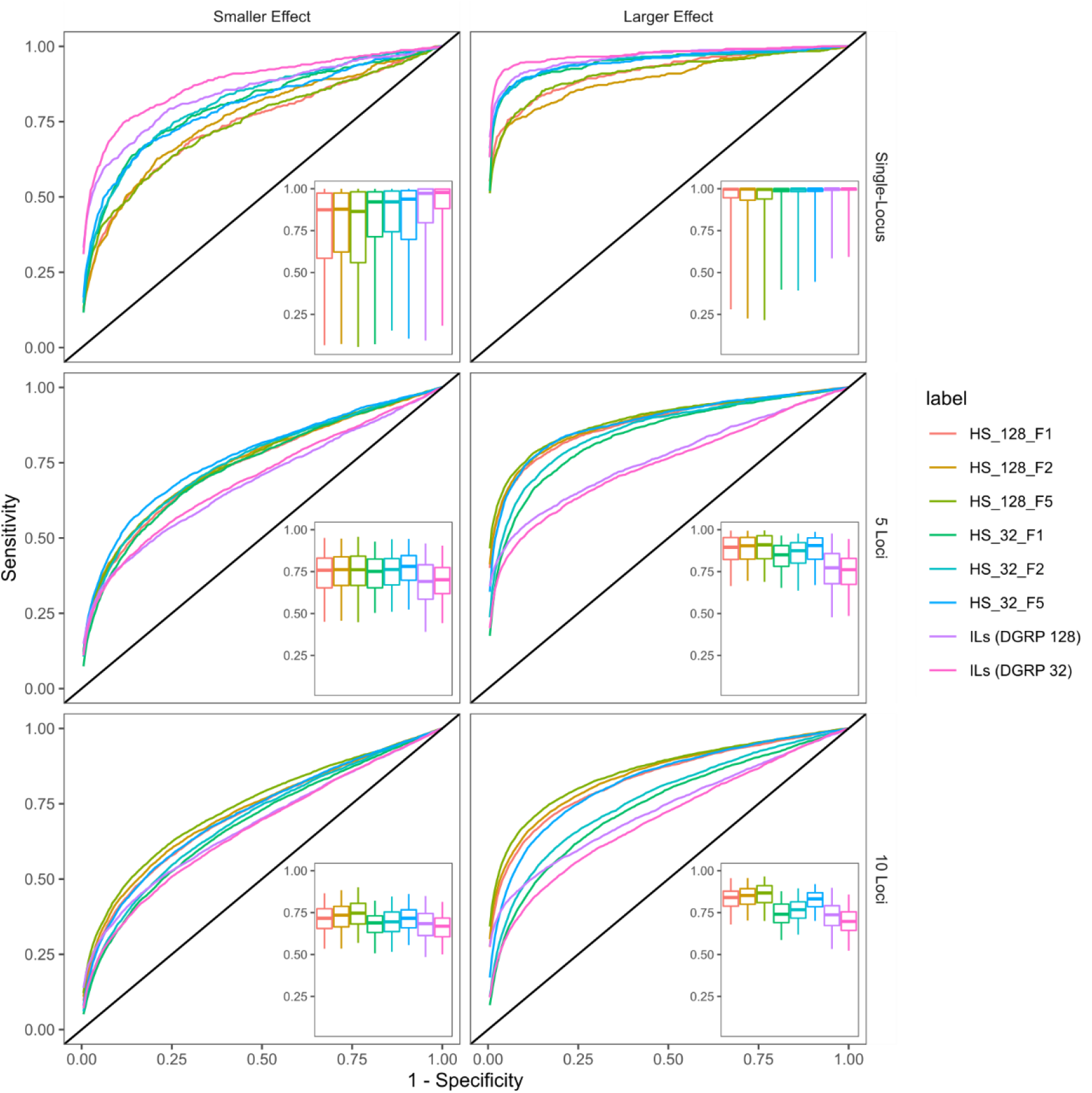
Receiver Operator Characteristic (ROC) curves for additional GWAS simulations. To generate single representative ROC curves across 500 GWAS simulations per population, we stepped through all specificity values and calculated the mean sensitivity. Inset boxplots display the median and interquartile range of Area Under the Curve (AUC) distributions, with whiskers spanning the middle 95% of data. **ILs**: inbred lines. **HS**: Hybrid Swarm populations founded by 32 or 128 lines.

**Supplemental Figure S13.**
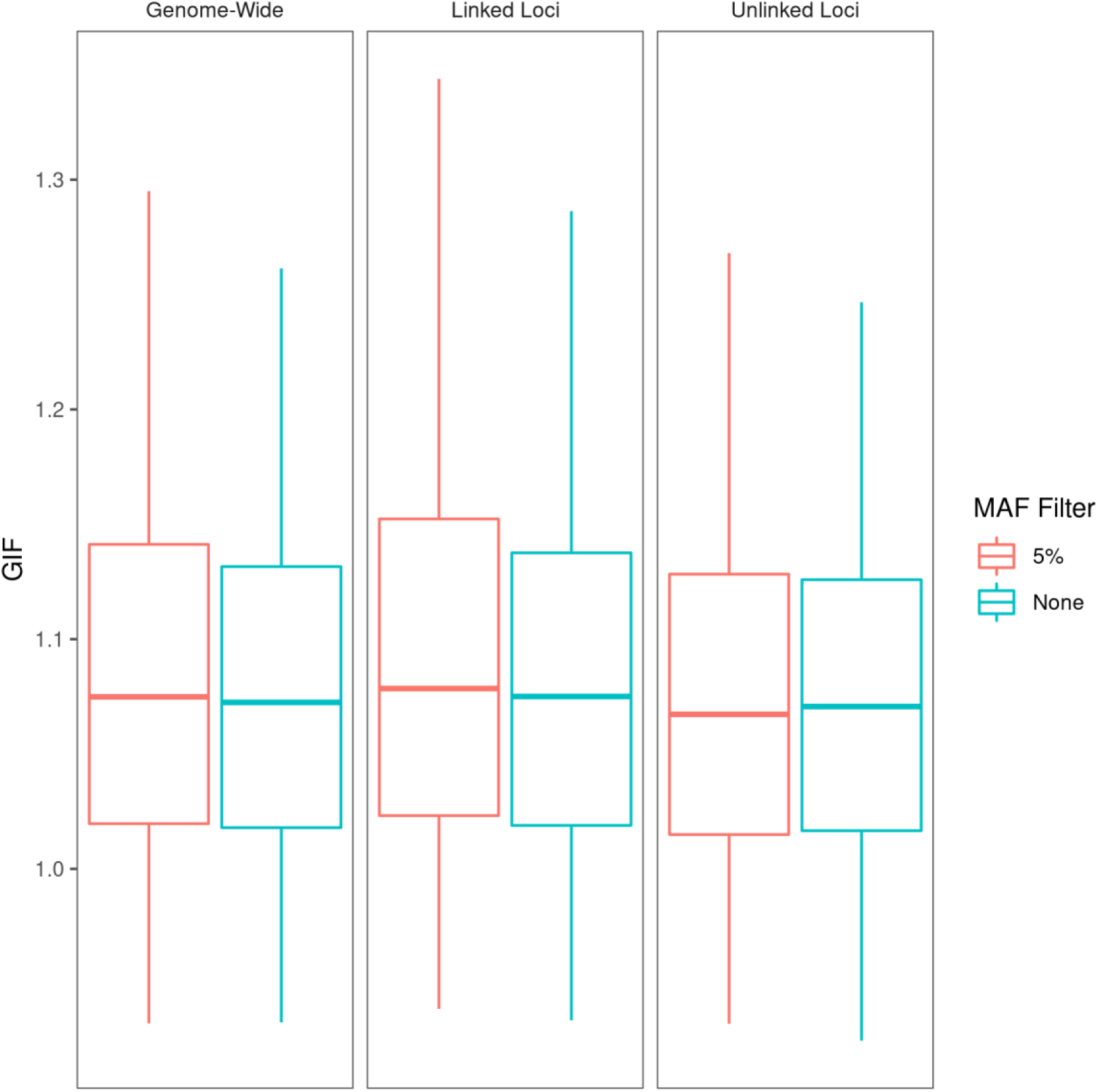
Genomic Inflation Factor with and without low-frequency filtering. Excluding alleles with a minor allele frequency below 5% does not reduce genomic inflation factor in inbred lines. Box depicts the middle 50% quantiles with whiskers extending to the middle 95% of data points. Traits were modeled as a single locus of large effect.

**Supplemental Table S1.** Accuracy of estimated number of recombination events following chromosome reconstruction. A high concordance correlation coefficient (Lin’s ρ) indicates agreement between estimated and true recombination counts for 400 reconstructed chromosomes (coalescent-derived populations) or chromosome arms (DGRP-derived populations). Coalescent-derived populations are described across a range of values for effective population size N_e_ and mutation rate μ. Δ represents the difference between estimated and true recombination counts, and σ_Δ_ represents the mean of 400 standard deviations of Δ. Reconstructions were performed with a maximum of 16 most-likely-ancestors with a HARP threshold of 0.99 (see methods for more details).

| Population | Coverage | N Founders | $N_e$ | $\mu$ | $\rho$ | Mean $\Delta$ | $\sigma_\Delta$ |
| --- | --- | --- | --- | --- | --- | --- | --- |
| DGRP | 0.005X | 128 | - | - | 0.201 | 3.06 | 3.11 |
| DGRP | 0.005X | 32 | - | - | 0.201 | 3.15 | 2.81 |
| DGRP | 0.05X | 128 | - | - | 0.986 | -0.015 | 0.25 |
| DGRP | 0.05X | 32 | - | - | 0.502 | 0.17 | 2.15 |
| Coalescent | 0.05X | 128 | $10^4$ | $1 \times 10^{-9}$ | 0.029 | -2.72 | 1.60 |
| Coalescent | 0.05X | 32 | $10^4$ | $1 \times 10^{-9}$ | 0.219 | -1.84 | 1.50 |
| Coalescent | 0.05X | 128 | $10^5$ | $1 \times 10^{-9}$ | 0.288 | -1.57 | 1.64 |
| Coalescent | 0.05X | 32 | $10^5$ | $1 \times 10^{-9}$ | 0.761 | -0.53 | 0.99 |
| Coalescent | 0.05X | 128 | $10^6$ | $1 \times 10^{-9}$ | 0.742 | -0.46 | 1.09 |
| Coalescent | 0.05X | 32 | $10^6$ | $1 \times 10^{-9}$ | 0.754 | -0.42 | 1.22 |
| Coalescent | 0.05X | 128 | $10^4$ | $5 \times 10^{-9}$ | 0.203 | -2.03 | 1.73 |
| Coalescent | 0.05X | 32 | $10^4$ | $5 \times 10^{-9}$ | 0.622 | -0.89 | 1.17 |
| Coalescent | 0.05X | 128 | $10^5$ | $5 \times 10^{-9}$ | 0.652 | -0.64 | 1.41 |
| Coalescent | 0.05X | 32 | $10^5$ | $5 \times 10^{-9}$ | 0.782 | -0.26 | 1.16 |
| Coalescent | 0.05X | 128 | $10^6$ | $5 \times 10^{-9}$ | 0.956 | -0.17 | 0.44 |
| Coalescent | 0.05X | 32 | $10^6$ | $5 \times 10^{-9}$ | 0.759 | -0.31 | 1.26 |
| Coalescent | 0.05X | 128 | $10^4$ | $1 \times 10^{-8}$ | 0.238 | -1.65 | 1.86 |
| Coalescent | 0.05X | 32 | $10^4$ | $1 \times 10^{-8}$ | 0.745 | -0.66 | 1.05 |
| Coalescent | 0.05X | 128 | $10^5$ | $1 \times 10^{-8}$ | 0.776 | -0.34 | 1.12 |
| Coalescent | 0.05X | 32 | $10^5$ | $1 \times 10^{-8}$ | 0.846 | -0.25 | 0.93 |
| Coalescent | 0.05X | 128 | $10^6$ | $1 \times 10^{-8}$ | 0.937 | -0.22 | 0.56 |
| Coalescent | 0.05X | 32 | $10^6$ | $1 \times 10^{-8}$ | 0.833 | -0.32 | 0.95 |

**Supplemental Table S2.** Test statistics and p-values for 270 pairwise comparisons of simulated GWAS AUC distributions. Our p-value significance threshold at α=0.05 is 1.85×10^-4^.

## Supplemental Methods

### Generating simulated reference panels across a range of diversity levels

To evaluate low-coverage reconstruction for various degrees of genetic diversity, we generated reference panels using haplotypes produced by coalescent models across a range of genetic diversity levels. Haplotypes were generated using the R (R Core Team 2016) package scrm (Staab *et al*. 2015) and subsequently restructured into VCF file format (Danecek *et al*. 2011). We generated ten independent panels for each of all 18 combinations of population size (N_e_=104, 105, 106), mutation rate (μ=10-9, 5×10-9, 10^-8^), and number of haplotypes (32, 128). The value θ for each simulation was defined as 4N_e_μ. We simulated a chromosome-length locus of 25 Mb with a recombination rate of 1.5 cM/Mb. SNP positions output by scrm (a decimal within the range of 0 to 1) were converted to base pair positions by multiplying the decimal by chromosome length (25 ×106 base pairs for our simulations) and rounding down to the nearest integer. Any sites with more than two alleles were converted to a biallelic site by discarding tertiary or quaternary alleles. Genotype values were re-coded as polarized signed integers: +1 for reference and −1 for alternate alleles. For every position, reference and alternate alleles were defined by randomly selecting one of the twelve non-repeating pairs of nucleotides. Reference genome FASTA files were created with a custom python script that generated a 25 million length string of nucleotide characters with weighted probability to achieve 45% GC-content, followed by replacing variable positions with their respective reference alleles.

### Extended details for simulating GWAS

#### Simulated haplotypes

Although the forward simulator we developed is efficient, it would not have been computationally feasible to simulate 500 fully independent mapping populations (per parameter combination) in a reasonable amount of time. Instead, we generated ten independent forward-simulated populations, and for each of those, generated fifty randomly permuted subsets. For a single simulated mapping population, we began by sampling (with replacement) a random subset of 5,000 individuals, out of 10,000 total individuals generated by forward-simulation. Then, we performed a permutation of haplotype ancestry with a new, randomly-ordered (equally sized) subset of founders. The permutation of ancestry was one-to-one, e.g. all haplotype blocks that were previously derived from founder X would be translated to founder Y, and blocks previously derived from Y would in turn be mapped to founder Z.

For all simulated GWAS, we began with a set of 129 DGRP haplotypes with the least missing data, e.g. high coverage and low levels of heterozygosity. This allowed us to perform leave-one-out subsampling for the 128-founder populations. In addition to Hybrid Swarm populations, which we ran through the simulated sequencing and mapping pipeline, we generated four additional types of mapping populations for comparing GWAS performance: Highly outbred (F50) populations; Inbred Lines (ILs) to represent the DGRP; and Recombinant Inbred Lines (RILs), similar to the Drosophila Synthetic Population Resource, or DSPR (King *et al*. 2012).

The F50 populations were generated with 128 founders in same manner as the Hybrid Swarm, except that populations were simulated over fifty non-overlapping generations of recombination instead of five generations. The ten resulting forward-simulated populations were resampled and permuted as we did with the Hybrid Swarms.

We simulated ten initial sets of 800 RILs using the same forward-simulator as previously described, each initialized with a random subset of eight DGRP haplotypes. Populations randomly recombined at a population size of 10,000 for fifty non-overlapping generations, after which 800 random male-female pairs of individuals were isogenzied through 25 generations of full-sibling mating. This scenario roughly corresponds to the DSPR. For computational simplicity, after the 25 generations of isogenization we removed any remaining residual heterozygosity by forcing the identity of a second chromosome copy to be identical to the first copy. We then sampled 5,000 draws (with replacement) of the 800 RILs followed by ancestry permutation as described above.

To simulate GWAS on Inbred Lines, no forward-simulation was necessary. For a single simulated population, we first randomly selected 128 DGRP lines, then randomly sample with replacement 5,000 times. As with hybrid swarm and RILs, for any parameter combination we generated a total of 500 mapping populations.

#### Liability model

Using the liability framework, we assign case or control phenotypes to mapping populations derived from DGRP chromosome 2L haplotypes. First, a genotype-dependent risk score is calculated for every individual, which is translated into probabilities for case or control group assignment. We parameterized our model such that individuals begin with equivalent odds of being assigned to case or control groups, and that probability is modified depending on allele status at causal loci.

For a given simulation, N causal loci are randomly selected with equal probability out of all segregating sites in the mapping population. The effect of individual causal loci are simulated as a Gaussian variable drawn with a small extent of noise (sd=0.005) with an expected (mean) value dependent on the allele frequency at that locus. Intermediate frequency loci (i.e. allele frequency = 0.5) are simulated to contribute on average no effect on case/control assignment; alleles near fixation decrease risk; rare alleles increase risk. The linear center of the curve, where most individual risk scores exist under the parameter combinations used in our simulations, approximates additive genetic architecture. The sigmoid tails allow for the continuous risk score to translate to a phenotype necessarily bounded by zero and one. The relationship between allele frequency, risk score, and phenotypic assignment is diagrammed in Supplemental Figure S9. Within the linear portion of the curve (which is the domain for the majority of simulations), a singleton minor allele will modify ‘case’ assignment probability by approximately +5% (small effect) or +10% (large effect), beginning at an initial assignment probability of 50%. The major allele, conversely, would decrease assignment by an opposite amount. An allele at 50% frequency will not modify case/control assignment (aside from a very small degree of random noise equal to about +/- 0.06%). Note that we performed GWAS simulations assuming 100% genotype accuracy.

We implemented a method of efficiently aggregating allele counts compatible with our haplotype map format. Briefly, haplotype map breakpoints across all individuals are sorted in ascending order. When iterating through ascending unique start and stop positions, between any pair of breakpoints, all SNPs will be comprised of the same number of each founding haplotype. Haplotype IDs could then be counted and sorted in the same column position order as the table containing polarized allele status (−1 for alternate, +1 for reference). Multiplying the genotype table by the haplotype count vector results in final allele counts, polarized negative for alternate alleles and positive for reference alleles.

## Notes

https://github.com/cory-weller/HS-reconstruction-gwas

### Competing Interest Statement

The authors have declared no competing interest.

### Summary of Updates

This version of the manuscript has been revised to make our simulation methodology more clear. This version also includes examples of our reconstruction pipeline on a handful of real-life hybrid swarm Drosophila melanogaster individuals.

